# Corticonigral projections recruit substantia nigra pars lateralis dopaminergic neurons for auditory threat memories

**DOI:** 10.1101/2024.11.04.621665

**Authors:** Lorenzo Sansalone, Rebekah C. Evans, Emily Twedell, Renshu Zhang, Zayd M. Khaliq

## Abstract

Dopaminergic neurons (DANs) in the lateral substantia nigra project to the tail of striatum (TS), which is involved in threat conditioning. Auditory cortex also contributes to threatening behaviors, but whether it directly interacts with midbrain DANs and how these interactions might influence threat conditioning remain unclear. Here, functional mapping revealed robust excitatory input from auditory and temporal association cortexes to substantia nigra pars lateralis (SNL) DANs, but not to pars compacta (SNc) DANs. SNL DANs exhibited unique firing patterns, with irregular pacemaking and higher maximal firing, reflecting different channel complements than SNc DANs. Behaviorally, inhibiting cortex to SNL projections impaired memory retrieval during auditory threat conditioning. Thus, we demonstrate robust corticonigral projections to SNL DANs, contrasting with previous observations of sparse cortical input to substantia nigra DANs. These findings distinguish SNL DANs from other nigral populations, highlighting their role in threatening behaviors and expanding knowledge of cortex to midbrain interactions.

## Introduction

Midbrain dopaminergic neurons (DANs), known to enable reward processing^1,2^, have been increasingly recognized for their role in aversive and threatening behaviors^3-6^. Dopamine signaling in regions associated with threat processing, such as the amygdala^7-9^, prefrontal cortex^10,11^, and striatum^12-15^, has been well documented. However, there has been comparatively less investigation into the involvement of specific dopaminergic neuron subpopulations and what circuit projections drive their activity during threatening behaviors.

Substantia nigra pars lateralis (SNL) DANs project to the tail of the striatum (TS), where they play a crucial role in processing salient, novel and aversive stimuli^16-21^, yet little is understood about their intrinsic properties and which synaptic circuits modulate their activity during threatening behaviors. Optogenetic stimulation of dopaminergic axons within the TS causes avoidance of novel objects^12^ while inhibition reduces auditory fear responses^15^. Chemical lesioning of these projections to TS reduces avoidance behaviors, suggesting their importance for threat avoidance^12^. Regarding projections regulating SNL DANs during aversive behaviors, the central nucleus of the amygdala (CeA) indirectly modulates SNL DANs by targeting SNL GABAergic neurons, which then disinhibit SNL DANs^22^. Moreover, anatomical studies using viral-genetic mapping have identified projections to substantia nigra DANs, showing that subthalamic nucleus (STN) provides particularly selective input to SNL DANs^16^. However, the excitatory projections that drive SNL DANs during threatening behaviors are not yet known.

Auditory threat conditioning^23^ engages the auditory cortex (AC), which then interacts with the amygdala^24^ and striatum^25,26^ to enable acquisition and establishment of auditory threat memories. However, whether cortical inputs engage DANs during threat signal processing through a direct corticonigral pathway remains unknown. Anatomical studies have identified cortical projections to substantia nigra DANs^16,27-29^ while functional results suggest that corticonigral projections to SNc DANs from motor cortex are sparse^30^. Whether AC provides direct projections to midbrain DANs located in SNc or SNL and whether these projections contribute to threat conditioning have not yet been determined.

Here, we demonstrate that the auditory association cortex (AAC), comprising secondary auditory cortex (AuV) and temporal association cortex (TeA), provides specific projections to SNL DANs that contribute significantly to memory retrieval during auditory threat conditioning. We show that optical activation of projections from the auditory cortex robustly increases firing in SNL DANs. By contrast, we found that lateral or medial SNc DANs (lSNc or mSNc) receive virtually no input from auditory association cortex. Comparative analysis of the firing properties of SNL, lateral SNc, and medial SNc DANs showed that SNL DANs represent an independent neuronal subpopulation characterized by irregular pacemaking and significantly higher maximal firing rates. These findings are further supported by biophysical experiments demonstrating differences in underlying ion channels expressed in SNL and SNc DANs. Finally, auditory threat conditioning experiments showed that disrupting synaptic transmission from SNL DANs to TS using virally-expressed tetanus toxin significantly reduced threat learning (CS-US association) during Pavlovian paradigms. Importantly, preventing synaptic release from auditory cortex to SNL DANs did not affect threat learning but greatly impaired the retrieval of auditory threat memories. Taken together, our findings reveal a corticonigral pathway that directly influences the activity of midbrain DANs involved in auditory-related aversive behaviors. These results offer new insight into how cortical-midbrain interactions influence dopaminergic transmission and contribute to threatening behaviors. These findings may enhance our understanding of the altered sensory processing associated with post-traumatic stress disorder (PTSD) and phobias^31^.

## Results

### SNL DA neurons exhibit distinct intrinsic firing properties relative to SNc DA neurons

Dopaminergic neurons (DANs) in both the substantia nigra pars lateralis (SNL) and pars compacta (SNc) neurons project to the tail of the striatum (TS), but whether SNL and SNc DANs differ in their morphology and physiology has not yet been determined. To identify SNL DANs, we first performed in situ hybridization experiments to detect cells co-expressing tyrosine hydroxylase (TH) with either calbindin or glutamate vesicular transporter 2 (VGluT2)^32-36^. Calbindin-positive (Calb+) and VGluT2-positive (VGlutT2+) DANs were present in the SNL and SNc (Extended Data Figure 1 and 4), consistent with previous findings^34^. Moreover, Calb+ DANs almost always co-expressed VGluT2 (Extended Data Figure 2). As an alternative strategy to identify SNL DANs, we employed an intersectional genetic approach by crossing either Calb-Cre or VGluT2-Cre mice to DAT-Flp::Ai65 mice (Extended Data Figure 3). This approach similarly revealed Calb+ and VGluT2+ DANs in both the SNL and SNc (Extended Data Figure 4), raising the question of whether these neurons comprise distinct subpopulations.

To compare DANs in the SNL and SNc, we first quantified the size of neurons by taking the cross-sectional somatic area of TH positive neurons using in situ hybridization in fixed midbrain slices (Figure 1a-c). SNc DANs had large cell bodies with no significant difference between medial SNc (mSNc) and lateral SNc (lSNc) neurons (somatic area; mSNc, 402.98 ± 11.75 μm^2^, n = 130; lSNc, 413.63 ± 20.28 μm^2^, n = 53, p = 0.78). However, we found that SNL neurons were significantly smaller (∼30%) with an average somatic area of 287.71 ± 12.72 μm^2^ (SNL: n = 47; SNL vs lSNc, p = 1.07 × 10^−5^; SNL vs mSNc, p = 2.61 × 10^−10^). Importantly, results from marker mouse lines were similar with the average somatic area of Calb+ and VGluT2+ SNL DANs being up to 60% smaller than SNc DANs (Extended Data Figure 4).

**Figure 1.**
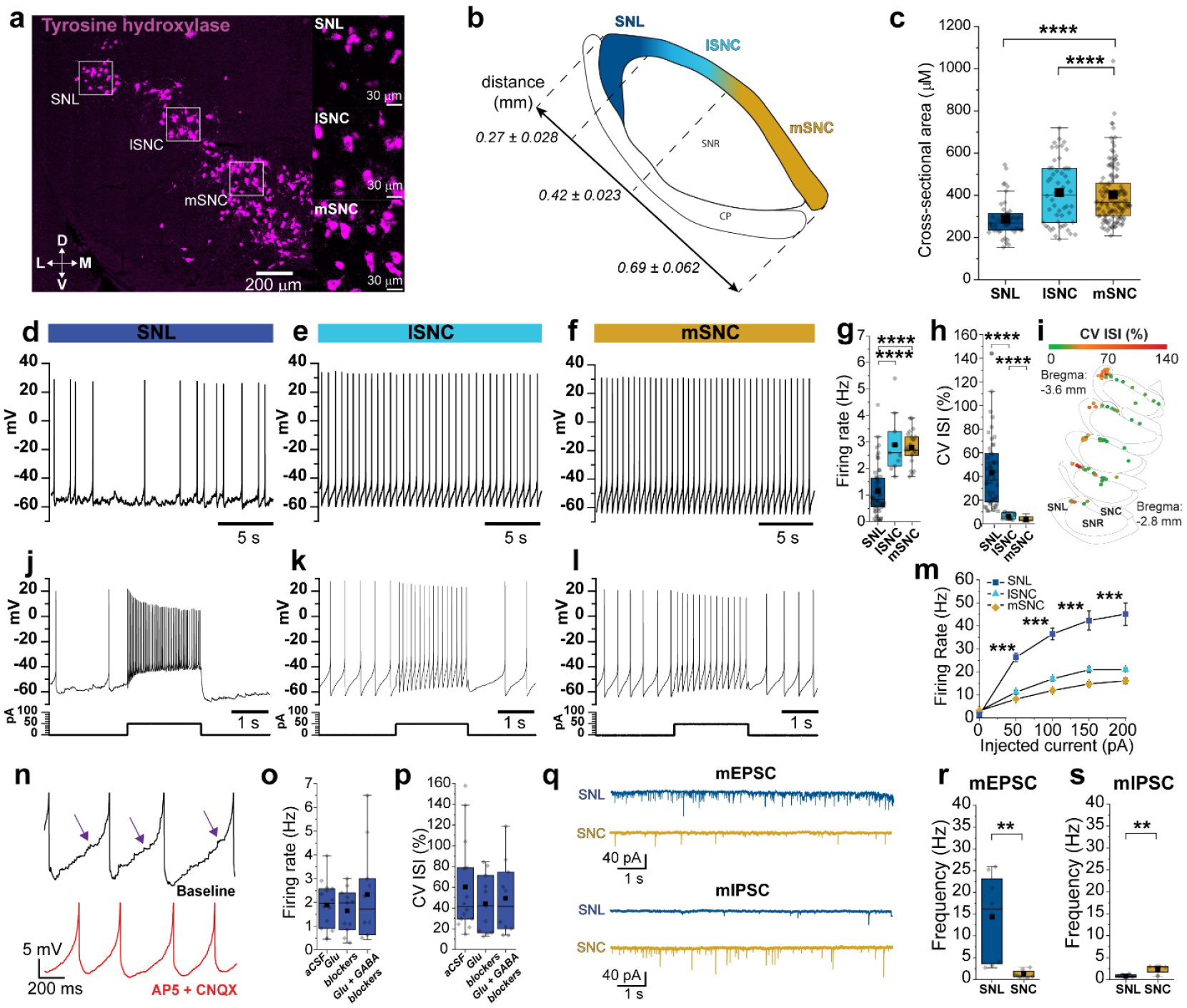
Distinct intrinsic firing and synaptic properties in SNL DA neurons. **a**, In situ hybridization of TH+ neurons. *Insets*, SNL (*top*), lateral SNc (*middle*), medial SNc (*bottom*). **b**, Illustrative map of nigral divisions. Numerical values represent average length of subdivision in mm ± s.e.m. **c**. Bar plots showing somatic cross-sectional areas from TH+ (SNL vs. lat SNc, p = 1.07 × 10^−5^; SNL vs med SNc, p = 2.61 × 10^−10^). **d-f**, Representative whole-cell recordings of DA neuron spontaneous firing. **g-h**, Firing rate and CV ISI averages in DANs. **I**, Plots of cell location within SN mediolateral axis with neurons color coded according to CV ISI. **J-L**, Representative recordings of evoked firing in DANs (same cells as above). **m**, Averaged frequency-current curves. **n**, *Top*, Representative trace of post-synaptic potentials during interspike voltage (*arrows*). *Bottom*, Trace from same neuron following AP5/CNQX. Note, suppression of PSPs. **o-p**, Firing rate and CV ISI averages from SNL DANs in control, plus glutamatergic receptor antagonists (AP5/NBQX), and plus GABA antagonists (AP5/NBQX/GBZ/CGP). **q**. Representative mEPSCs (*top*) and mIPSC (*bottom*) from SNL and SNC DANs recorded in the presence of TTX plus gabazine (mEPSC) or AP5-NBQX (mIPSC). **r, s**, Averaged mEPSCs and mIPSC frequencies for SNL vs SNc DANs. Box whiskers represent 25-75% percentiles, solid squares are mean value, horizontal box lines represent medians. **\****p <* 0.05, **\*\****p <* 0.01, **\*\*\****p <* 0.001, **\*\*\*\****p <* 0.0001

We next performed whole-cell patch-clamp electrophysiology experiments to examine the intrinsic firing properties of SNL DANs. We found that SNL DANs fire spontaneously at significantly lower rates compared to SNc DANs (Figure 1d-g) (avg firing rate; SNL, 1.15 ± 0.13 Hz, n = 53; mSNc, 2.80 ± 0.13 Hz, n = 23; lSNc, 2.90 ± 0.32 Hz, n = 11; SNL vs mSNc, p = 1.08 × 10^−10^; SNL vs lSNc, p = 2.07 × 10^−6^). Pacemaking in SNc DANs was highly rhythmic^37^. By contrast, we found that pacemaking in SNL DANs is highly irregular, with an average coefficient of variation of interspike interval (CV ISI) of 43.14 ± 4.26 % (n = 48) compared to SNc DANs which display CV ISI lower than 10% (Figure 1h; CV ISI; mSNc, 3.94 ± 0.42 %, n = 23; lSNc, 6.37 ± 0.75 %, n =11; SNL vs mSNc, p = 7.55 × 10^−19^; SNL vs lSNc, p = 7.15 × 10^−12^).

Similar observations were made with cell-attached and perforated-patch electrophysiology experiments, suggesting that the irregular firing in SNL DANs did not reflect wash-out of intracellular signaling proteins in whole-cell recordings (Extended Data Figure 5). Moreover, CV ISI increased across the mediolateral axis of the substantia nigra (Figure 1i), from mSNc to SNL. Additionally, we found an inverse relationship between CV ISI and the capacitance of DANs (Extended Data Figure 7), indicating that the observation of higher CV ISI correlates with smaller neuron sizes. Comparison of firing in VGluT2+ DANs located in SNL revealed that these differ substantially from those located in lSNc (Extended Data Figure 6). Testing maximal firing rates with direct current-injections (Figure 1j-m) revealed that SNc neurons exhibited low maximal firing rates (average maximal firing rate, mSNc, 20.93 ± 0.78 Hz, n = 13; lSNc, 16.12 ± 1.04 Hz, n = 4) consistent with past work^38^. By contrast, we found that SNL DANs exhibited substantially higher maximal firing rates of 45.08 ± 4.85 Hz (n = 15; SNL vs mSNc, p = 5.34 × 10^−4^; SNL vs lSNc, p = 5.16 × 10^−4^). Thus, SNL and SNc DANs intrinsic firing properties suggest that they represent two different dopaminergic subpopulations.

When examining the interspike voltage in SNL DANs whole-cell patch-clamp recordings, we noticed the presence of subthreshold post-synaptic potentials (PSPs). The bulk of these PSPs were abolished by CNQX and D-AP5, inhibitors of excitatory synaptic AMPA and NMDA receptors (Figure 1n and Extended Data Figure 10). Importantly, inhibition of synaptic transmission had no significant effect on the firing rates (Figure 1o; control vs. AP5/NBQX, p = 0.65; control vs. AP5/NBQX/GBZ/CGP, p = 0.89) or CV ISI (Figure 1p, control vs. AP5/NBQX, p = 0.43; SNL AP5/NBQX/GBZ/CGP, p = 0.44), suggesting that the irregular pacemaking results from the intrinsic properties of SNL DANs. We found that SNL DANs have more depolarized AHP voltages during spontaneous pacemaking compared to SNc DANs (Extended Data Figures 8a,b), which is further supported by the presence of lower SK-conductances (Extended Data Figures 8c,d). We also recorded smaller sag voltages during direct hyperpolarization in SNL DANs relative to SNc DANs (Extended Data Figures 9a,b), indicative of weaker HCN recruitment (Extended Data Figures 9c-e). To more accurately quantify synaptic transmission, we performed voltage-clamp recordings and compared miniature excitatory and inhibitory post-synaptic currents (mEPSC and mIPSC) in SNL and SNc DANs (Figure 1q). We found that SNL DANs exhibit substantially higher mEPSC frequency with an average of 14.32 ± 3.47 Hz compared to 1.26 ± 0.31 Hz for SNc neurons (Figure 1r, SNL: n = 8; SNc, n = 8, p = 3.11 × 10^−4^). By contrast, the frequency of mIPSCs was significantly lower in SNL DANs compared to SNc DANs (Figure 1s; SNL, 0.82 ± 0.14 Hz, n = 6; SNc, 2.37 ± 0.34 Hz, n = 8, p = 0.012). Therefore, our analysis of miniature events shows that SNL DANs exhibit a high ratio of excitatory to inhibitory (E/I) events, suggesting that they receive primarily excitatory input, which differs substantially from SNc DANs which are known to be governed mainly by inhibition^39^.

### Functional input mapping reveals exclusive innervation of SNL DA neurons by auditory association cortex

To examine the major excitatory inputs to SNL DANs, we first retrogradely labeled projections to the SNL by injecting Cholera Toxin Subunit B conjugated to Alexa Fluor 647 (CTB647) into the SNL (Figure 2a). We found strong retrograde labeling in layers 2/3 and 5 of the auditory association cortex (AuV/TeA) (Figure 2a,b and Extended Data Figure 12), a high-order auditory processing station that integrates auditory information with experience-dependent cues^40^.

**Figure 2.**
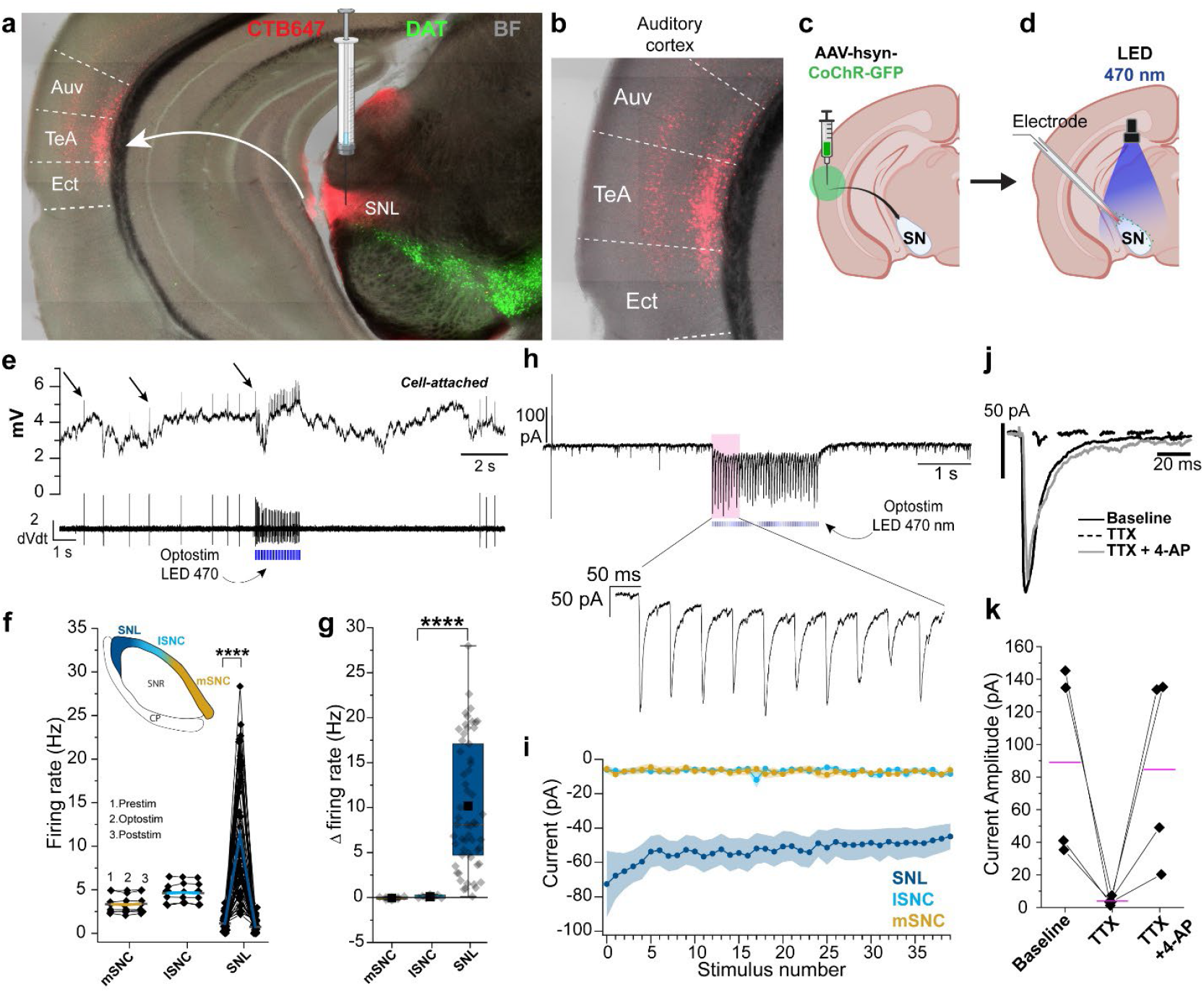
Auditory association cortex projects specifically to SNL DA neurons. **a**, Confocal image of coronal slice from DAT-Cre Ai9 mouse showing DANs (*green*) and retrogradely labeled neurons in cortex from CTB-647 injection in SNL (*red*). **b**, Magnification of auditory cortex with retrogradely labeled neurons in ectorhinal cortex (EcT), temporal association cortex (TeA) and ventral auditory cortex (AuV) **c**, Illustration showing location for stereotaxic injection with AAV-CoChR in AuV/TeA (*green*) (Created with BioRender.com). **d**, Illustration showing optogenetic stimulation of AuV/TeA terminals while recording with patch-pipette from substantia nigra DANs (Created with BioRender.com). **e**, *Top*, Cell-attached recording from SNL DAN during optical stimulation (470 nm, 2 s, 20 Hz, 2 ms), *arrows* indicating action potentials. *Bottom*, 1^st^ derivative of the trace above showing clear resolution of action potentials. **f**, Firing rates before (1), during (2) and after (3) optical stimulation for mSNc, lSNc and SNL DANs. *Black dot*, average firing rate from 5 sweeps from single neuron. *Colored lines*, means. **g**, Graph showing absolute % change (Δ) in firing rate during optical stimulation. Box whiskers represent 25-75% percentiles. **h**, *Top*, representative voltage clamp recording of SNL DAN responding to optical stimulation with clear oPSC. *Bottom*, magnification of evoked currents. **i**, Averaged oPSCs amplitude for mSNc, lSNc and SNL DANs. Shaded areas represent ± s.e.m. **j**, SNL DANs oPSC from single light pulse (2 ms, 470 nm) for baseline (solid black), TTX (dotted black) and TTX + 4-AP (solid grey). **k**, Averaged oPSCs amplitude for SNL DANs for baseline, TTX and TTX + 4-AP. Purple lines represent means. **\***p *<* 0.05, **\*\***p *<* 0.01, **\*\*\***p *<* 0.001, **\*\*\*\***p *<* 0.0001.

To functionally examine the impact of these cortical projections, we performed stereotaxic viral injections of AAV-CoChR in AuV/TeA of DAT-Cre Ai9 mice (Figure 2c) and tested the impact of optical stimulation of AuV/TeA projections while performing cell-attached recordings in SNL, lSNc and mSNc DANs (Figure 2d). Optical stimulation of AuV/TeA inputs (2 ms of 470 nm light pulses delivered at 20 Hz for 2 s) dramatically increased the firing rate of SNL DANs (Figure 2e) from 1.14 ± 0.11 Hz pre-stimulation to 11.32 ± 0.93 Hz (Figure 2f; n = 57, p *=* 2.38 × 10^−15^) corresponding to an average increase of 10.18 Hz ± 0.94 Hz (Figure 2g). Surprisingly, SNc DANs did not respond significantly to optical stimulation with mSNc average firing rate going from a baseline of 3.35 ± 0.38 Hz to 3.29 ± 0.37 Hz during optical stimulation and lSNc DANs average firing rate going from a baseline of 4.61 ± 0.44 Hz to 4.70 ± 0.42 Hz during optical stimulation (Figure 2j; mSNC, n = 8, p = 0.36; lSNc: n = 7, p = 0.23) which corresponds to a change in firing rate of -0.05 ± 0.05 Hz and 0.09 ± 0.07 Hz, respectively (Figure 2g).

We confirmed these findings by testing the presence of synaptic currents (oPSCs) in response to optical stimulation of the AuV/TeA in the same neurons (Figure 2h). Importantly, SNL DANs consistently displayed oPSCs while mSNc and lSNc DANs did not show clear currents upon stimulation (Figure 2i). When performing voltage-clamp recordings, in the presence of tetrodotoxin (TTX) and 4-aminopyridine (4-AP), SNL DANs consistently responded to a single light pulse with a clear oPSC (Figure 2j,k; average oPSC amplitude, n = 4, 89.11 ± 29.50 pA) demonstrating that projections from AuV/TeA to SNL DANs are monosynaptic in nature.

### STN and PPN differentially innervate SN DANs

Next, we evaluated retrograde labeling from SNL CTB647 injections (Figure 2a) in the two main excitatory nuclei providing input to midbrain DANs, the pedunculopontine nucleus (PPN) and subthalamic nucleus (STN)^27^. While we found labeling in both STN and PPN (Figure 3a and 3e), we noticed that most of the retrogradely labeled cells of the STN were located in the most dorsolateral region (Figure 3a). To confirm this, we have performed injected CTB555 and CTB647 in the SNL and lSNc of a C57WT mouse. We found that most retrogradely labeled neurons projecting to lSNc were located in the center of the STN while retrogradely labeled neurons projecting to SNL were, once again, located in the dorsolateral region of the STN (Extended Data Figure 11). These findings suggest that SNL and SNc DANs are likely to receive innervation from distinct neurons within the STN.

**Figure 3.**
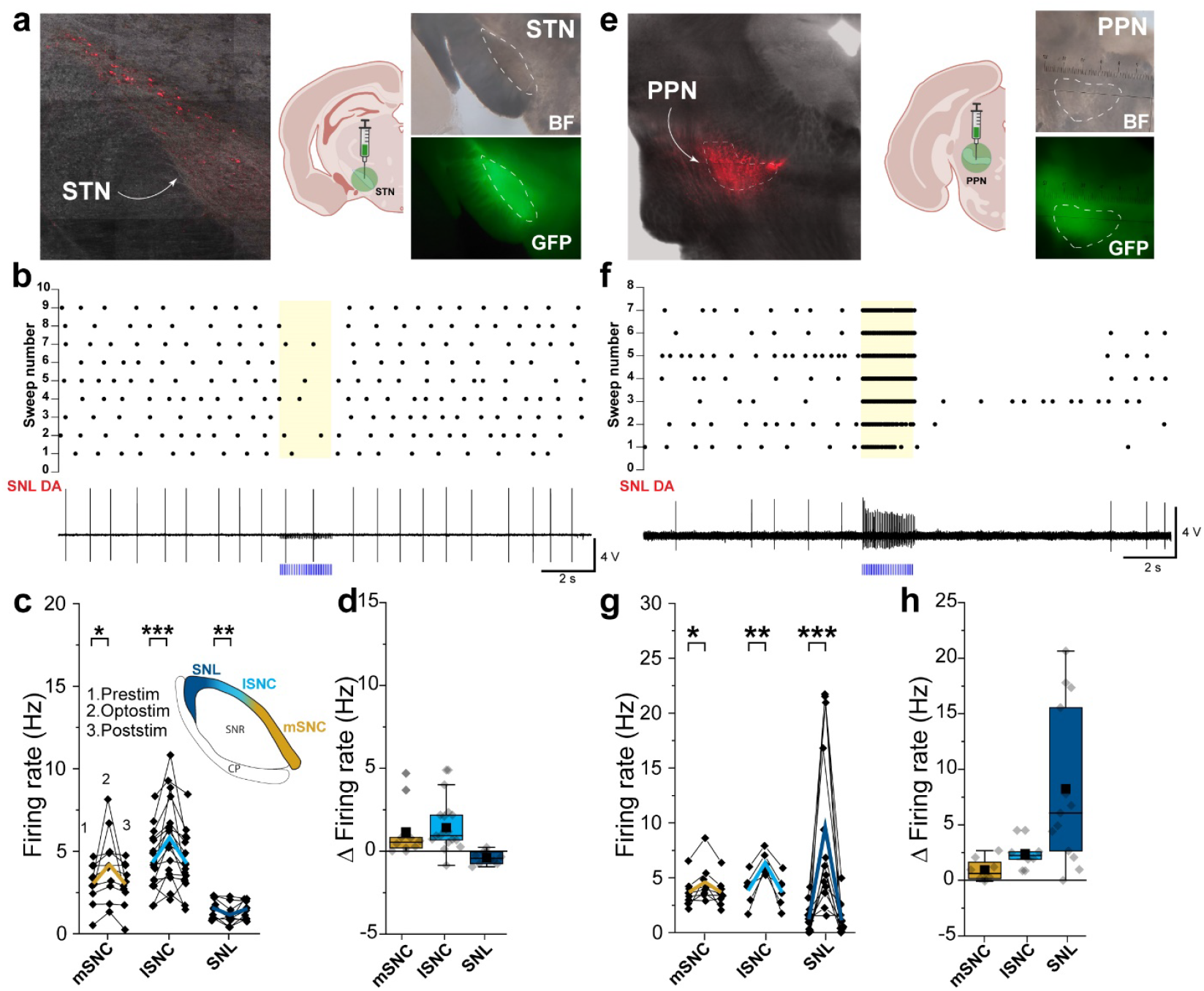
STN and PPN projections to SNL DA neurons. **a**, *Left*, confocal image showing retrogradely labeled neurons in STN from CTB647 injections in SNL in a DAT-Cre Ai9 mouse. Note that most neurons are located withing the dorsolateral section of STN. *Center*, Cartoon showing location of stereotaxic injection with AAV-CoChR (Created with BioRender.com). *Right*, Brightfield and fluorescence confocal images showing the extent of STN targeting with AAV-CoChR. **b**, Raster plot of individual stimulation trials (*top*) and representative 1^st^ derivative of current-clamp recording (*bottom*) showing SNL DAN response to optical stimulation (470 nm, 2 s, 20 Hz, 2 ms) of STN terminals infected with CoChR. **c**, Firing rates before (1), during (2) and after (3) optical stimulation for mSNc, lSNc and SNL DANs. Black dot, average firing rate from 5 sweeps from single neuron. Colored lines, means. **d**, Graph showing absolute % change (Δ) in firing rate during optical stimulation of STN terminals. Box whiskers represent 25-75% percentiles. **e**, *Left*, confocal image showing retrogradely labeled neurons in PPN from CTB647 injections in SNL in a DAT-Cre Ai9 mouse. *Center*, Cartoon showing location of stereotaxic injection with AAV-CoChR (Created with BioRender.com). *Right*, Brightfield and fluorescence confocal images showing the extent of PPN targeting with AAV-CoChR. **f**, Raster plot of individual stimulation trials (*top*) and representative 1^st^ derivative of current-clamp recording (*bottom*) showing SNL DAN response to optical stimulation (470 nm, 2 s, 2 ms) of PPN terminals infected with CoChR. **g**, Firing rates before (1), during (2) and after (3) optical stimulation for mSNc, lSNc and SNL DANs. Black dot, average firing rate from 5 sweeps from single neuron. Colored lines, means. **h**, Graph showing absolute % change (Δ) in firing rate during optical stimulation of STN terminals. Box whiskers represent 25-75% percentiles. **\***p *<* 0.05, **\*\***p *<* 0.01, **\*\*\***p *<* 0.001, **\*\*\*\***p *<* 0.0001.

To functionally examine the impact of STN and PPN projections to substantia nigra DAN subpopulations, we first performed stereotaxic viral injections of AAV-CoChR in STN and PPN of DAT-Cre Ai9 mice (Figure 2a, *right*; figure 2e, *right*) and tested the effects of optogenetic stimulation on DANs in SNL and SNc. Following optical stimulation of STN projections, we found that SNc DANs consistently increased their firing suggesting that SNc neuron are innervated as previously reported^41-43^, with a significant change in firing rate from a pre-stimulation baseline of 3.04 ± 0.39 Hz to 4.21 ± 0.66 Hz for mSNc, and from 4.41 ± 0.43 Hz to 5.83 ± 0.60 Hz for lSNc (Figure 3c; mSNc, n = 10, p *=* 0.026; lSNc, n = 17, p = 0.0004), which corresponds to an average firing rate increase of 48 ± 19.88 % and 36.38 ± 9.99 %, respectively (Figure 3d). SNL DANs did not respond consistently to optical stimulation of STN terminals as shown in the raster plot from independent trials reported in Figure 3b. Upon stimulation, we recorded a small change in firing rate from 1.54 ± 0.18 Hz to 1.15 ± 0.20 Hz (Figure 3c; SNL, n = 10, p *=* 0.0044) which corresponds to an absolute firing rate decrease of 25.58 ± 9.24 % (Figure 3d). Next, we tested PPN projections. We found that SNL DANs strongly respond to optical stimulation of PPN terminals as shown in the raster plot reported in Figure 3f. We found that both SNc sections responded to optical stimulation of PPN projections, as previously reported^44^, with mSNc and lSNc DANs showing firing increments from 3.68 ± 0.47 Hz to 4.61 ± 0.64 Hz and from 3.97 ± 0.60 Hz to 6.36 ± 0.40 Hz, respectively (Figure 3g; mSNC, n = 8, p *=* 0.017; lSNc, n = 6, p *=* 0.002). These firing increments correspond to absolute firing rate increase of 29.24 ± 14.41 % and 84.40 ± 37.63 for mSNc and lSNc, respectively (Figure 3h). Importantly, we recorded a larger increase in firing rate for SNL DANs from 1.37 ± 0.34 Hz to 9.60 ± 2.14 Hz (Δ = 3153.42 ± 1911.65 %) (Figure 3h; SNL, n = 13, p *=* 6.40 × 10^−4^; and Figure 3h). Altogether, our examinations of the intrinsic firing and synaptic inputs of DA subpopulations demonstrate that SNL DANs are a functionally distinct subpopulation of neurons.

### DA projections to the tail of the striatum contribute to auditory threat learning

SNL DANs contribute to aversive learning and their projections to TS are involved in learning of threat avoidance^12,15,20,21^. Therefore, we next evaluated the role of DA released in TS during auditory threat conditioning^23^. To test this, we inhibited synaptic transmission from SNL projection terminals by expressing tetanus toxin light chain (Tet-LC or TetTx) in DANs by using DAT-Cre mice injected in TS with AAV-DIO-eGFP-TetTx or AAV-DIO-eGFP (Figure 4a, *left*) and confirmed viral expression 4 weeks later (Figure 4A, *right*). Our threat conditioning paradigm (Figure 4b) consisted of two phases on day 1 - a habituation phase during which pure tones (5 kHz, 30s duration; conditioned stimulus, CS) were presented alone for five times at random intervals, followed by a conditioning phase during which the presented pure tones co-terminated with an electric footshock (0.6 mA, 1 s; unconditioned stimulus, US). After 24 h, on day 2, the conditioned animals were placed in a novel context (Context B, Figure 4b) and underwent a retrieval phase during which pure tones were presented without footshock to test threat memory expression. For each phase, animal freezing was quantified as absolute % freeze during pure tone presentation. During the habituation phase, mice did not freeze in response to pure tones (Figure 4c) (habituation phase, % freeze; control, 0.19 ± 0.20%, N = 8; TetTx, 0.41 ± 0.27%, N = 8; p = 0.73). During the conditioning phase, however, both groups displayed increasing freezing behavior with the TetTx-treated mice freezing significantly less than control (Figure 4c,e) (conditioning phase, % freeze; control, 23.21 ± 5.49 %, N = 8; TetTx, 7.78 ± 3.46 %, N = 8; p = 0.048). During the retrieval phase, control mice continued to freeze in response to the CS tone while TetTx treated mice froze significantly less (Figure 4d,e) (retrieval phase, % freeze; control, 47.13 ± 5.63 %, N = 8; TetTx, 13.32 ± 3.37 %, N = 8; p = 0.001). Therefore, our results demonstrate that the inhibition of neurotransmitter release from SNL DANs in TS interferes with the encoding and consolidation of auditory threat memory. These results align with the view that DA in TS plays a role in promoting defensive (innate) reactions to threatening stimuli^12,15,20-22^.

**Figure 4.**
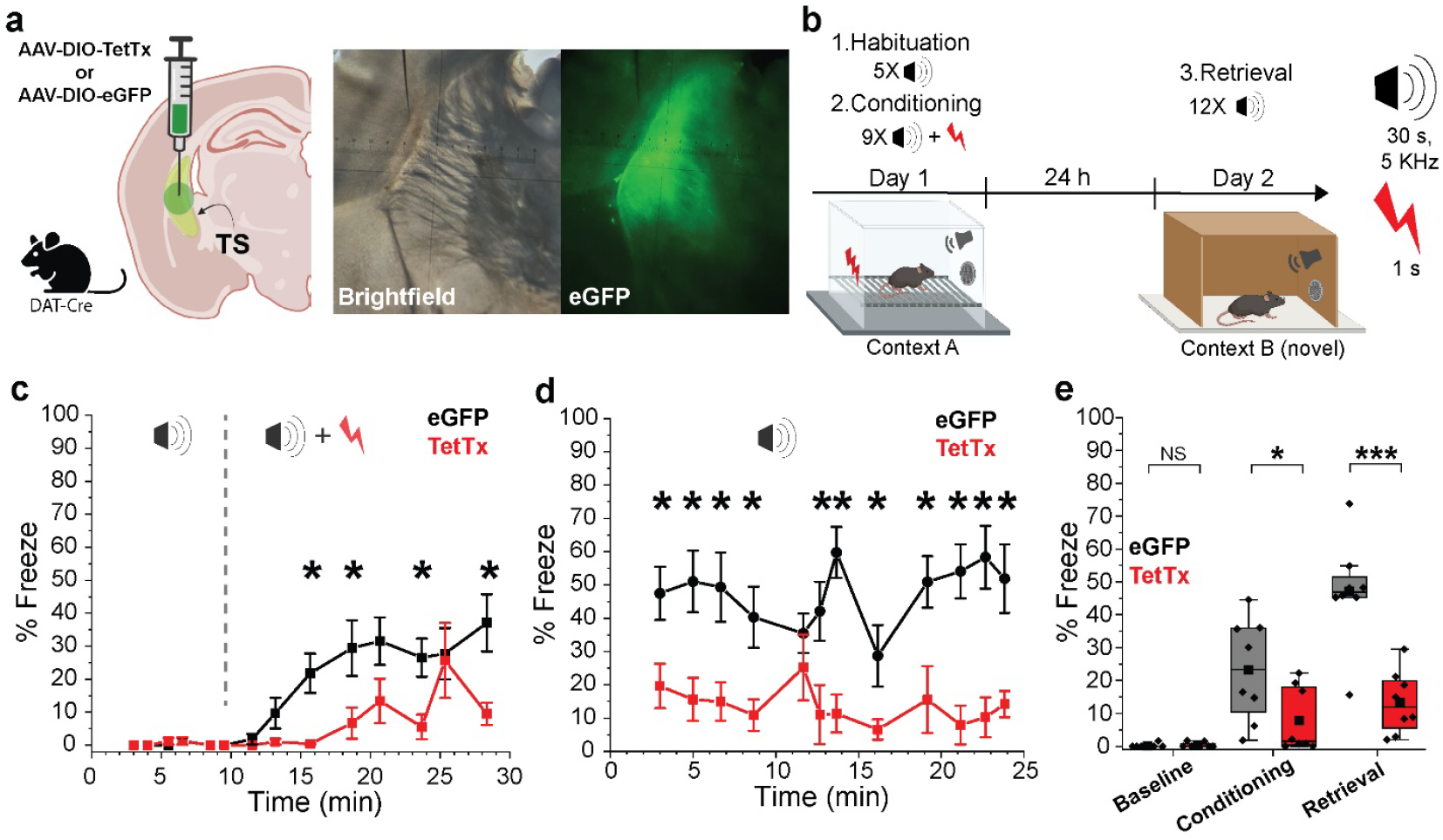
Tail of the striatum (TS) projecting DA neurons contribute CS-US association during auditory threat conditioning. **a**, *Left*, Illustration showing TS injection site for AAV-DIO-TetTx-eGFP or AAV-DIO-eGFP (Created with BioRender.com). *Right*, Representative brightfield and fluorescence images showing the extent of viral infection in TS. **b**, Behavioral paradigm used for auditory threat conditioning. **c**, Graph showing binned % freeze during auditory tones for the habituation phase (before dotted line) and conditioning phase (after dotted line). Each squared symbol represents the average for 30 s pure tone (N = 8 for each group). Bars are ± s.e.m. **d**, Graph showing absolute % freeze during auditory tones for the retrieval phase. Each squared symbol represents the average for a 30 s pure tone (N = 8 for each group). Bars are ± s.e.m. **e**, Averaged % freeze during baseline, conditioning, and retrieval phases for control (eGFP) and TetTx treated mice.

### Cortical transmission to SNL DA neurons is required for retrieval of auditory threat memories

Our results showed that auditory association cortex (AuV/TeA) provides strong excitation to SNL DANs, however the behavioral consequences of cortical input to SNL DANs are unknown. To test the contribution of AuV/TeA->SNL projections in threat learning, we performed a procedure involving two stereotaxic injections in C57WT mice - the first consisted of AAV9-Cre injected into the SNL to retrogradely target AuV/TeA neurons, followed by a second injection of AAV-DIO-eGFP-TetTx into the AuV/TeA to selectively express TetTx in SNL-projecting neurons (Figure 5a,b). Next, we performed Pavlovian threat conditioning over three days using the same context (A) during habituation, conditioning, and retrieval (Figure 5c). We used a train of frequency modulated (FM) tones (5-20 kHz, 0.5 s tones at 1 Hz for 30 s duration; conditioned stimulus, CS) to assess the function of AuV/TeA->SNL projections because it was shown that this cortical area is important for processing complex (FM) sounds^26,45^. We did not find significant differences in freezing behavior during the habituation and conditioning phases for control and treated mice, however, a clear trend towards diminished freezing behavior during conditioning was observed for TetTx-treated mice (Figure 5d, 5e and 5g; % freeze; habituation phase - control, 7.04 ± 1.48 %, N = 8; treated, 3.63 ± 2.05 %, N = 6; p *=* 0.13; conditioning phase - control, 26.13 ± 5.42 %, N = 8; treated, 11.42 ± 4.66 %, N = 6; p *=* 0.11). During the retrieval phase, we recorded a significant increase in % freeze for control vs treated mice (Figure 5f and 5g; % freeze; retrieval phase - control, 31.90 ± 5.54 %, N = 8; treated, 6.48 ± 2.65 %, N = 6; p = 0.0047). To confirm this effect was not influenced by contextual freezing (location-induced) but actually due to auditory threat memory, we quantified animal freezing before exposing the animals to FM sweeps during the conditioning and retrieval phases (Figure 5h). There was no significant difference between the two groups (Figure 5h; contextual % freeze; pre-conditioning phase – control, 3.34 ± 1.12 %, N = 8; treated, 0.825 ± 0.825 %, N = 6; p = 0.061; pre-retrieval phase - control, 24.59 ± 5.69 %, N = 8; treated, 27.27 ± 8.85 %, N = 6; p = 0.061; p = 0.70) suggesting contextual freezing was experienced equally by both groups.

**Figure 5.**
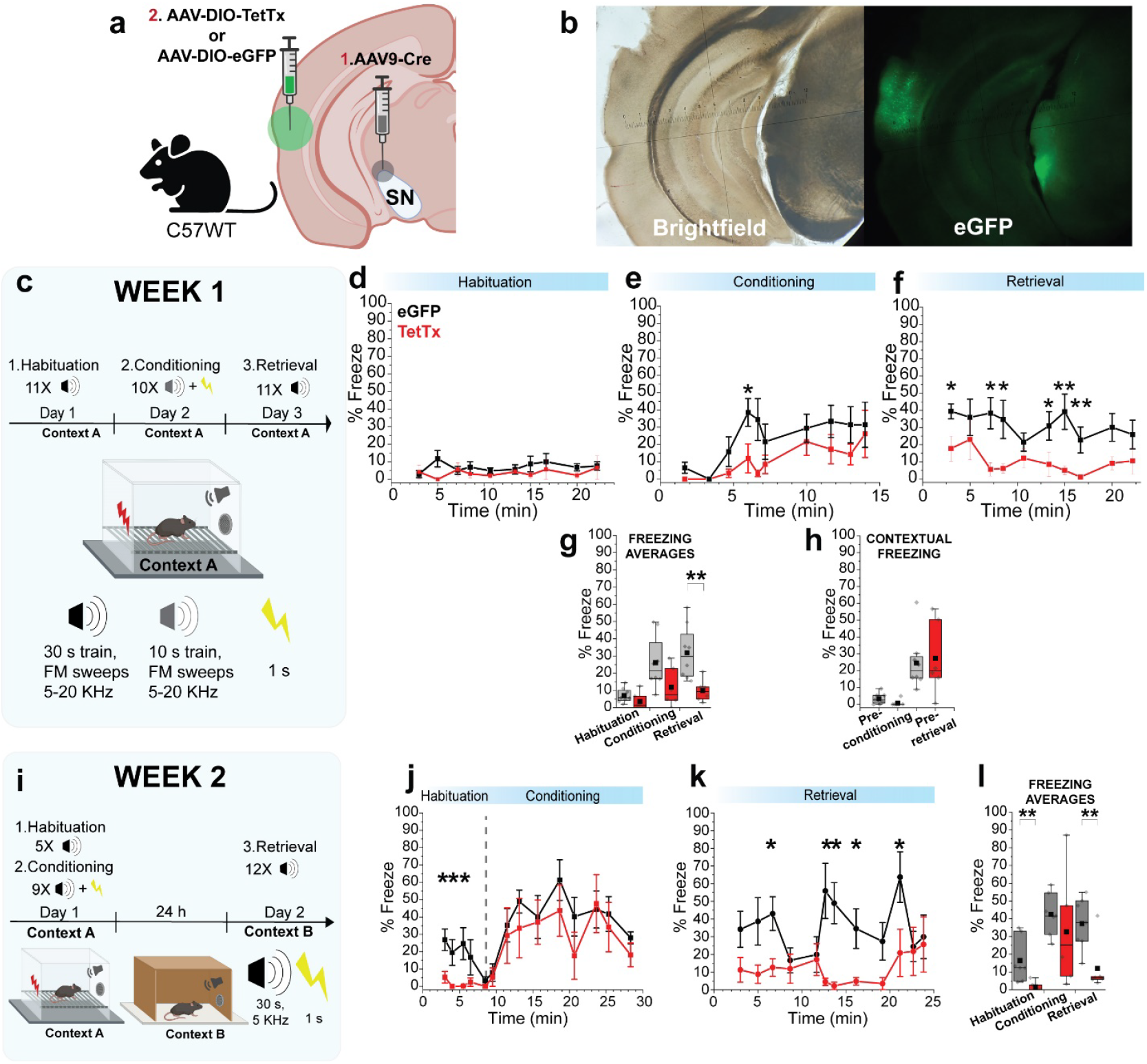
Auditory threat conditioning is modulated by auditory association cortex neurons projecting to SNL. **a**, Illustration showing infection strategy with 2 viral injections: AAV9-Cre in SNL followed by AAV-DIO-eGFP or AAV-DIO-eGFP-TetTx in AuV/TeA (Created with BioRender.com). **b**, Representative brightfield and fluorescence images showing the extent of viral infection in AuV/TeA/SNL. **c**, Behavioral paradigm used for auditory threat conditioning on week 1. **d-f**, Graphs showing binned % freeze for habituation, conditioning, and retrieval phases. Squared symbols represent % average freeze during FM tone trains (control, N = 8; TetTx, N = 6). **g**, Averaged % freeze during FM tones. Square symbols represent averages from a single mouse. **h**, Averaged % contextual freeze before conditioning and retrieval phases, respectively. Measurements were performed before the first tone was played and while the mouse was exploring the environment. Squared symbols represent a single mouse. **i**, Behavioral paradigm used for auditory threat conditioning on week 2. **j-k**, Graph showing binned % freeze during habituation/conditioning, and retrieval phases. Squared symbols represent the freeze average during a single pure tone (control, N = 8; TetTx, N = 6). **l**, Averaged % freeze during auditory tones. Every squared symbol represents average from a single mouse. All bars shown with ± s.e.m.

To evaluate if the conditioned mice had the ability to store the auditory threat memory for longer periods, after 1 week, we reconditioned our groups using the same protocol from Figure 4, which uses pure tones and different contexts for habituation/conditioning and retrieval phases (Figure 5i). During the habituation phase, upon presentation of pure tones, we found a stark difference between groups, with control mice displaying clear freezing behavior compared to treated mice which did not freeze (Figure 5j, *before dotted line*, and 5l; % Freeze; habituation phase – control, 16.44 ± 4.73 %, N = 8; treated, 1.58 ± 1.13 %, N = 6; p = 0.005). This indicates that treated mice were not able to store auditory threat memories from the previous trials, performed 7 days before. During the conditioning phase, however, we noticed both groups responded with comparable freezing behavior (Figure 5i, *after dotted line*, and 5l; conditioning phase – control, 42.48 ± 4.47 %, N = 8; treated, 32.66 ± 12.76 %, N = 6; p = 0.37). This confirms that TetTx-treated mice can still process sounds and associate them with threatening/aversive stimuli (CS-US association). Importantly, we noticed a significant difference during the retrieval phase when treated mice displayed diminished freezing behavior compared to control during CS recall (Figure 5k and 5l; % freeze; retrieval phase - control, 37.28 ± 5.49 %, N = 8; treated, 12.02 ± 5.95 %, N = 6; p = 0.01). Once again, these findings suggest that synaptic inactivation of AuV/TeA-SNL projections significantly impairs the long-term establishment of auditory threat memory, while still allowing the real-time processing of CS-US association for threatening stimuli.

## Discussion

We demonstrate that SNL DANs comprise a functionally distinct subpopulation defined by their intrinsic properties, synaptic inputs and behavioral contribution to threat conditioning. Comparison of the intrinsic properties of substantia nigra DANs show that SNL DANs are distinguished by irregular firing and high maximal firing rates which likely results from different ion channels complement compared to SNc DANs. Examination of synaptic projections from different brain nuclei show that SNL DANs are distinct, receiving little and distinct STN input relative SNc DANs (Figure 3A and Extended Data Figure 11). As the major difference in projection patterns between DAN subpopulations, we reveal that the auditory association cortex [comprised of secondary auditory cortex (AuV) and temporal association cortex (TeA)], Extended Data Figure 12) provides robust and specific input to SNL DANs which are likely to contribute to threat conditioning. By contrast, our results show that SNc DANs are not targeted by cortical inputs. Importantly, we show that abolishing synaptic release in auditory association cortex projecting to SNL does not affect threat learning (CS-US association) but greatly impairs the retrieval of auditory threat memory. Thus, our results expand the knowledge on the dopaminergic system in the modulation of animal behavior, demonstrating that aversive responses and threat conditioning likely involve direct communication from cortical projections to DANs in the SNL.

Although the firing properties of DAN subpopulations within the SNc have been well described^46-49^, there has been almost no exploration of the intrinsic firing properties of SNL DANs. In this study, we extensively characterized the firing properties of SNL DANs. We found that SNL DANs fire rapidly at substantially higher maximal firing rates relative to canonical SNc DANs which exhibit low maximal rates as a result of their tendency to enter depolarization block^50^. In addition, we found that spontaneous firing in SNL DANs is irregular, consistent with in vivo extracellular recordings of midbrain DANs^51^. The persistence of irregular firing in SNL DANs, even in the presence of synaptic blockers (Figure 1O), demonstrates that this is an intrinsic membrane property of these neurons. Moreover, we show that SNL DANs are smaller in size, have higher maximal firing rate, and display lower levels of hyperpolarization-activated cyclic nucleotide-gated (HCN) and small-conductance calcium-activated potassium (SK) conductances compared to SNc DANs. Importantly, lower expression of SK channels in SNL DANs correlates with irregular firing as observed in VTA DANs^52-54^. These findings show that SNL DANs are clearly distinguished from other substantia nigra DANs.

The observed differences in the morphology and physiological properties of SNL and lSNc DANs that we report here have implications for understanding the role of dopamine in the tail of the striatum (TS). The intrinsic properties strongly determine the cell’s input-output relationship. Thus, our observations raise the possibility that the SNL and lSNc may provide distinct signals to TS, as DANs in both regions project to this area. Moreover, this distinct signaling may also apply to the subset of VGluT2 DANs which we found exhibiting differences between SNL and SNc DANs (Extended Data Figure 6). Therefore, in future behavioral studies that use VGluT2-or Calb-Cre mouse lines to study dopamine release in TS, it would be interesting to examine separately the contributions of SNL and lateral SNc neurons to threatening behaviors.

Analysis of miniature post-synaptic currents (minis, excitatory: mEPSCs, inhibitory: mIPSCs) revealed that SNL DANs exhibit a high-frequency of mEPSCs compared to mIPSCs, suggesting that they receive predominantly excitatory input. This surprising result stands in contrast to our previous knowledge of canonical SNc DANs that have been shown to receive primarily inhibitory synaptic input^39^. Moreover, this finding supports the idea that DANs in the SNL and SNc participate in different circuits, but also raises the question of what projections may provide unique inputs to SNL DANs. Previous anatomical studies using retrograde viral tracing showed that TS-projecting DANs are distinguished by receiving a strong excitatory projection from the subthalamic nucleus (STN)^16,27^. Additionally, previous work based on direct electrical stimulation of STN in anesthetized rats showed that dopamine is exclusively released in TS but not in the dorsolateral striatum (DS)^55^, which is contrast with previous literature showing that high-frequency stimulation of STN induces striatal dopamine release in rats and pigs^56,57^. Importantly, TS-projecting neurons originate from both the SNL and the neighboring lateral SNc which presents a challenge in distinguishing them by retrograde tracing. In our functional synaptic experiments, we found that activation of STN projections did not result in strong excitation of SNL DANs – most cells had no response in their firing. However, we found that activation of STN projections increased firing in SNc DANs (Figure 3c). These data were consistent with retrograde tracing showing that two different STN neuron populations are labeled following cholera toxin subunit B (CTB) injections into SNL and SNc (Figure 3 and Supplementary Figure 11). These results demonstrate the importance of using anatomical approaches along with functional electrophysiological experiments when evaluating connectivity of neighboring neuronal populations such as SNL and SNc DANs.

Importantly, our study revealed robust corticonigral projections from the auditory association cortex to SNL DANs, making these neurons unique among midbrain DANs. Following the earliest proposals of a corticonigral pathway from a century ago, there have been many studies that have tested for its existence, but the results were unclear. The most compelling evidence has been provided by anatomical studies that report only sparse connections from cortex to SNc DANs^27,58^. Functional evidence for corticonigral projections has been very limited and often ambiguous, particularly in *in vivo* experiments, due to possible indirect excitation through STN. A recent study tested motor cortex (M1 and M2) inputs to SNc DANs and found that optogenetic activation of M1/M2 projections resulted in measurable responses in a small number of SNc DANs tested (3 out of 35 DANs)^30^. By contrast, our findings here demonstrate that the vast majority of SNL DANs exhibit strong increases in firing upon optical stimulation of auditory association cortex projections compared to SNc DANs, which did not show any functional responses. We demonstrate that SNL DANs represent the only substantia nigra region receiving strong monosynaptic input from auditory association cortex (Extended Data Figure 12). Thus, corticonigral input likely contributes to the higher frequency of mEPSCs that were observed in SNL compared to SNc DANs.

Our data show that tetanus toxin inhibition of synaptic transmission from AuV/TeA to SNL significantly reduces animal freezing during the retrieval phase of a threat conditioning paradigm, suggesting that this corticonigral pathway is important for threat memory expression. This result is supported by previous work showing that cortical neurons located in AuV/TeA are critical for memory acquisition and expression^26,59^. Interestingly, it has been shown that the primary auditory cortex responds to frequency modulated (FM) sweeps only, while AuV/TeA responds to both FM sweeps and pure tones^26^, thus supporting our results (Figure 5).

Furthermore, AuV/TeA neurons that send projections to the lateral amygdala were found to be responsive to FM sweeps but not pure tones, which suggests that AuV/TeA neurons involved in pure tone conditioning are likely to use different output pathways. Our SNL-targeted retrograde tracing experiments with cholera toxin subunit B (CTB) led to the identification of labeled cortical neurons in layers 2/3 and 5 of AuV/TeA. This result, together with our behavioral findings, suggests that the same AuV/TeA neurons described in previous reports may participate in pure tone conditioning by exclusively recruiting SNL DANs, while FM responsive AuV/TeA neurons may project to both SNL and lateral amygdala. Finally, our data is further supported by previous work showing that cortical neurons located in layers 2, 3 and 5 of AuV/TeA exhibited activity dependent changes in excitatory genes in mice that underwent threat conditioning paradigms^59^.

Our data may have implications for understanding human psychiatric disorders such as post-traumatic stress disorder (PTSD) and phobia where misevaluation of threat value of sensory stimuli may trigger abnormal physiological reactions. The severity of clinical symptoms in PTSD and phobia has been linked to sensory areas^31^, suggesting that sensory processing dysfunction might contribute to symptoms’ severity. Importantly, the unique properties of SNL DANs, including faster firing and higher input resistance, may contribute to rapid sensory responses that may be advantageous for threat learning. Here, we report that the interaction of auditory association cortex and dopaminergic neurons in the SNL contributes to threat conditioning, raising the possibility that this pathway may be involved in circuit dysfunction in PTSD and phobia. Altogether, our findings provide a new framework aimed at defining the mechanisms underlying cortical sensory processing to downstream areas during aversive behaviors.

## Supporting information

Sansalone et al 2024_Supplementary Material

## Methods

### Ethical compliance

All procedures were conducted in accordance with the guidelines established by the Animal Care and Use Committee for the National Institute of Neurological Disorders and Stroke (NINDS) and the National Institutes of Health (NIH).

### Animals

Experiments were carried out using male and female mice at 4-24 weeks of age. The following strains were used: DAT^IRESCre^ (B6.SJL-Slc6a3^tm1.1(cre)Bkmn^/J), The Jackson Laboratory, Strain #: 006660); VGluT2^IRESCre^ (Slc17a6^tm2(cre)Lowl^/J), The Jackson Laboratory, Strain #: 016963); Calb1-^IRES2-Cre-D^ (B6;129S-Calb1^tm2.1(cre)Hze^/J), The Jackson Laboratory, Strain #: 028532); DAT-Flp (Slc6a3^em1(flpo)Hbat^/J), The Jackson Laboratory, Strain #: 035436) which was obtained from Helen Bateup lab); Ai65(RCFL-tdT)-D (B6.129S-Gt(ROSA)26Sor^tm65(CAG-tdTomato)Hze^/J), The Jackson Laboratory, Strain #: 021875); Ai9(RCL-tdT) (B6.129S6-Gt(ROSA)26Sor^tm9(CAG-tdTomato)Hze^/J), The Jackson Laboratory, Strain #: 007905); C57BL/6 (C57BL/6NCrl, Charles River Laboratories, Strain #: 027); Tyrosine hydroxylase-GFP (Th-GFP; C57BL/6 background^3860^).

### Stereotaxic surgery

All stereotaxic injections were conducted on a Stoelting QSI (Cat# 53311). Mice were maintained under anesthesia for the duration of the injection with 1.5% isoflurane and allowed to recover from anesthesia on a warmed pad. At the end of the injection, the needle was raised 1-2 mm for a 10-minute duration before needle was removed. All mice were allowed to recover in their cage after injections and received subcutaneous ketoprofen (10 mg/kg) for 3 consecutive days post-injection. Mice were used for ex vivo electrophysiology or behavioral experiments 3-5 weeks after injections. The retrograde tracer cholera-toxin subunit B conjugated to Alexa fluor™ 647 (CTB-647, ThermoFisher scientific Cat # C34778) was bilaterally injected into the SNL (ML: ± 2.0, AP: - 3.0, DV: - 4.1) of C57WT or DAT-Cre Ai9 mice. AAV-CoChR (AAV1-hSyn-CoChR-GFP, UNC vector core, Boyden) was injected bilaterally into AuV/TeA (ML: ± 4.5, AP: - 3.0, DV: - 3.4), STN (ML: ± 1.6, AP: - 1.5, DV: - 4.9) and PPN (ML: ± 1.1, AP: - 4.4, DV: - 3.7) of DAT-Cre Ai9 mice.

AAV-DIO-eGFP-Tet-LC (AAVDJ-CMV-DIO-eGFP-2A-TeNT, Stanford University Gene Vector and Virus Core, Cat #GVVC-AAV-71) and AAV-DIO-eGFP (AAVDJ-CMV-DIO-eGFP, Stanford University Gene Vector and Virus Core, Cat #GVVC-AAV-12) were injected bilaterally into TS (ML: ± 3.4, AP: - 0.7, DV: - 3.0) and AuV/TeA (ML: ± 4.5, AP: - 3.0, DV: - 3.4) of C57WT mice.

AAV9-Cre (AAV0-hSyn-Cre-WPRE-hGH, Addgene Cat # 105553) was injected bilaterally into the SNL (ML: ± 2.0, AP: - 3.0, DV: - 4.1) of C57WT mice.

### Slice preparation

Mice were anesthetized with isoflurane, decapitated, and brains extracted. Coronal midbrain slices (200 μm) were prepared using a vibratome (Leica VT1200S). Slices were cut in ice-cold, oxygenated, slicing solution containing the following (in millimolar (mM)): 198 glycerol, 2.5 KCl, 1.2 NaH2PO4, 20 HEPES, 25 NaHCO3,10 glucose, 10 MgCl2, 0.5 CaCl2, 5 Na-ascorbate, 3 Na-pyruvate, and 2 thiourea. Slices were then incubated for 30 min at 34 °C in oxygenated holding solution containing the following (in millimolar (mM)): 92 NaCl, 30 NaHCO3, 1.2 NaH2PO4, 2.5 KCl, 35 glucose, 20 HEPES, 2 MgCl2, 2 CaCl2, 5 Na-ascorbate, 3 Na-pyruvate, and 2 thiourea.

Slices were then stored in same holding solution at 20-25 °C, with constant carbogen perfusion, and electrophysiological recordings were performed within 1 h to 6 h.

### Electrophysiological recordings

Slices were continuously superfused at 2.7 ml/min with warm (34°C), oxygenated extracellular aCSF recording solution containing the following (in millimolar (mM)): 125 NaCl, 25 NaHCO3, 1.25 NaH2PO4, 3.5 KCl, 10 glucose, 1 MgCl2, and 2 CaCl2 (Osmolarity: 290-310 mOsm). Neurons were visualized with a 60x objective using a BX61WI Olympus microscope equipped with a Hamamatsu digital camera ORCA-ER (C4742-80). Recordings were obtained using low-resistance pipettes (2.2 - 5 MΩ) pulled from filamented borosilicate glass (World Precision Instruments, Cat #1B150F-4) with a flaming/brown micropipette puller (Sutter Instruments, Model P-97). Cell-attached and whole-cell current-clamp recordings were made using borosilicate pipettes filled with internal solution containing (in mM) 122 KMeSO3, 9 NaCl, 1.8 MgCl2, 4 Mg-ATP, 0.3 Na-GTP, 14 phosphocreatine, 9 HEPES, 0.45 EGTA, 0.09 CaCl2 (Osmolarity: 280 mOsm). Some experiments included 0.1% - 0.3% neurobiotin (Vector Laboratories, Inc., Cat # SP-1120) in the internal solution for post hoc visualization. Whole-cell voltage-clamp recordings were made using borosilicate pipettes filled with internal solution containing (in mM) 120 CsMeSO3, 20 Tetraethylammonium chloride, 2 MgCl2, 4 Mg-ATP, 0.3 Na-GTP, 14 phosphocreatine, 10 HEPES, 10 EGTA, 2 QX314, 0.03 ZD7288 (Osmolarity: 280 mOsm) and cells were held at - 70 mV for AMPA/GABA currents and +40 mV for NMDA currents. For H-currents recordings, voltage-clamp experiments were made using borosilicate pipettes filled with internal solution containing (in mM) 130 KMeSO3, 30 Tetraethylammonium chloride, 10 NaCl, 2 MgCl2, 4 Mg-ATP, 0.3 Na-GTP, 14 phosphocreatine, 10 HEPES, 10 BAPTA-K (Osmolarity: 280 mOsm). Access resistance was monitored and recordings with R_a_ > 25 MΩ were discarded. Liquid junction potential (−8 mV) was not corrected. All experiments were conducted between 33-36 °C.

### Optogenetics experiments

Whole-field optogenetic activation of CoChR infected axons in brain slice was achieved by either a white LED (Prizmatix) sent through a FITC filter (HQ-FITC; U-N41001; C27045) or a blue (470nm) LED (Thorlabs, LED4D067) sent to the tissue via a silver mirror or through the FITC filter. Light intensity measured at the objective back aperture ranged from 1 – 25 mW. Light activation was given as a single pulse lasting 2 ms or as a 2 s, 20 Hz train with 2 ms pulses, unless otherwise specified.

### Reagents

Patch-clamp recordings and optogenetics experiments, where indicated, were performed in the presence of one or more of the following drugs: 20 mM 2,3-Dioxo-6-nitro-7-sulfamoyl-benzo[f]quinoxaline (NBQX) or 6-Cyano-7-nitroquinoxaline-2,3-dione (CNQX) to block AMPA receptors, 50 mM (2R)-Amino-5-phosphonopentanoate (D-AP-5) to block NMDA receptors, 50 mM Picrotoxin (PTX) or 10 μM Gabazine (GBZ) to block GABA_A_ receptor, 1 mM Tetrodotoxin (TTX) to block voltage-gated sodium channels, 200 μM 4-Aminopyridine (4-AP) to block voltage-gated potassium channels, 500 nM CGP-55845 to block GABA_B_ receptors, 200 nM Apamine to block SK channels. Glucose, glycerol, and salts used to make slicing, holding and perfusing aCSF solutions were purchased from Sigma-Aldrich. Drugs were purchased from Tocris Bioscience and Sigma-Aldrich. All drugs were reconstituted as indicate by the manufacturer and prepared as aliquots in deionized water or DMSO and stored at -20 °C or at -80 °C.

### Data analysis

Signals were digitized with a Digidata 1590 interface, amplified by a Multiclamp 700B amplifier, and acquired using pClamp 13 software (Molecular Devices). Data were sampled at 50 kHz and filtered at 10 kHz (or 2 KHz for analysis of mEPSC frequency). Data were analyzed using custom codes procedures in IgorPro (WaveMetrics). All recordings were performed in DANs. DANs were targeted by their anatomic location, size, and presence of the fluorescence reporter tdTomato. Analysis of electrophysiological data was conducted in Igor (Wavemetrics) and Clampfit. mEPSC or mIPSC were analyzed using Easy Electrophysiology. Unless otherwise specified, Mann-Whitney U tests (unpaired) or t-tests were used to compare two groups. For repeated comparisons, paired-t tests determined significance of the dataset. Data in text is reported as Mean ± SEM and error bars are ± SEM unless otherwise specified. Boxplots show medians, 25^th^ and 75th (boxes) and outliers 1.5 IQR (whiskers) percentiles. Biological replicates include samples from at least 3 separate mice. Exact P values are provided in the texts or figure legends.

### Immunohistochemistry, clearing, confocal imaging

After electrophysiology, slices were fixed for 2 – 12 h in 4% Paraformaldehyde (PFA) in phosphate buffer (PB, 0.1M, pH 7.6). Slices were then rinsed and stored in PB until immunostaining. For the immunostaining/CUBIC clearing, all steps are performed at room temperature on shaker plate. Slices (200 μm) were placed in CUBIC reagent 1 for 1 day, washed in PB 3x 1 hour each, placed in blocking solution (0.5% fish gelatin (sigma) in PB) for 3 hours. Slices were then directly placed in primary antibodies (sheep anti-TH and/or streptavidin Cy5 conjugate and/or rat anti-mCherry) in PB at a concentration of 1:1000 for 2-3 days. Slices were washed 3 times for 2 hours each and placed in secondary antibodies (Alexa 568 anti-sheep, or Alexa 488 anti-rat at 1:1000 in PB) for 2 days. After PB washed 3 times for 2 hours each, slices were placed in CUBIC reagent 2 overnight. Slices were mounted on slides in reagent 2 in frame-seal incubation chambers (Bio-Rad SLF0601) with coverslips. Slices were imaged on a Zeiss LSM 800 using Zen Blue software.

### Fluorescence in situ hybridization (FISH) - RNAscope^®^

*In situ* hybridization was performed on 16 μm thick midbrain slices from a fresh-frozen mouse brain cut on a cryostat (Leica CM3050 S). All FISH reagents used are commercially available from ACD bio, and procedures for the Multi-Plex FISH process were followed as recommended on ACDbio.com. Channels used for this study were TH, DAT (Slc6a3), Calb1 and VGluT2 (Slc17a6). Slices were imaged on a Zeiss LSM 800 using Zen Blue software.

### Auditory threat conditioning and analysis

Pavlovian auditory threat conditioning in Figure 4 consisted of one continuous habituation- conditioning session (Session 1) occurring in Context A followed by one retrieval (or recall) session, 24 h later, occurring in Context B (Session 2). Context A was a cage equipped with a speaker and floor-placed parallel metal rods that can deliver foot shocks. Context B was a cage equipped with a speaker and a plastic panel (white) on top of the metal rods as well as black and white design panels on the side walls. Context B was cleaned with acetic acid before mice placement. Between sessions, each chamber was cleaned with 70% ethanol solution. During session 1, 5 tones (CS, 5 kHz, 30 s, 75-80 dB) were delivered randomly with intertone intervals of 60 - 180 s, after which 8 tones (5 kHz, 30 s, 75-80 dB) co-terminating with a foot shock (US, 0.6 mA, 1 s) were randomly delivered. During session 2, 11 tones (5 kHz, 30 s, 75-80 dB) were randomly delivered with intertone intervals of 60 - 180 s. Experiments and data analysis were carried out using Freezeframe (Actimetrics). The percentage of time spent freezing (% Freezing) was calculated using Freezeframe software and represents the average time the animal freeze during the tones (CS) duration. Freezing threshold were manually determined based on software recommendation to prevent quantification of simple resting-pause positions instead of real freezing behavior. Pavlovian auditory threat conditioning in Figure 5 was performed over 2 weeks. For week 1, habituation, conditioning and retrieval were performed separately on consecutive days, respectively day 1, day 2, and day 3. For habituation and retrieval, 11 frequency modulated (FM) sound trains made of 0.5 s tones delivered at 1 Hz (5-20 KHz, 30 s, 75-80 dB) with random intertrain intervals of 60 - 180 s. For conditioning, 10 FM sound trains made of 0.5 s tones delivered at 1 Hz (5-20 KHz, 30 s, 75-80 dB) with random intertrain intervals of 60 - 180 s. For week 2, the same paradigm used for Figure 4 was followed.

## Authors contribution

L.S. and Z.M.K. conceived the project, designed experiments, and interpreted the data. L.S. wrote the first draft of the manuscript. L.S. and Z.M.K edited and wrote the manuscript. L.S., E.T. and R.C.E. performed experiments. L.S., E.T. R.C.E. and Z.M.K analyzed the data. R.Z. and L.S. performed stereotaxic surgeries.

## Acknowledgements

We thank the members of the Khaliq laboratory for their insightful discussions and comments on this manuscript. This study was funded by a fellowship to L.S. from the Center for Compulsive Behaviors (CCB), at the NIH Intramural Research Program. This work was supported by an NINDS Intramural Research Program Grant NS003135 to Z.M.K. Funding for this work was also provided by Aligning Science Across Parkinson’s (ASAP-020529) to Z.M.K. through the Michael J. Fox Foundation for Parkinson’s Research (MJFF).

## References

1 Bromberg-Martin, E. S., Matsumoto, M. & Hikosaka, O. Dopamine in Motivational Control: Rewarding, Aversive, and Alerting. Neuron 68, 815–834 (2010).

2 Bech, P. et al. Striatal Dopamine Signals and Reward Learning. Function 4, zqad056 (2023).

3 Fadok, J. P., Dickerson, T. M. K. & Palmiter, R. D. Dopamine Is Necessary for Cue-Dependent Fear Conditioning. The Journal of Neuroscience 29, 11089 (2009).

4 Pezze, M. A. & Feldon, J. Mesolimbic dopaminergic pathways in fear conditioning. Progress in Neurobiology 74, 301–320 (2004).

5 Zafiri, D. & Duvarci, S. Dopaminergic circuits underlying associative aversive learning. Frontiers in Behavioral Neuroscience 16 (2022).

6 Hamati, R., Ahrens, J., Shvetz, C., Holahan, M. R. & Tuominen, L. 65 years of research on dopamine’s role in classical fear conditioning and extinction: A systematic review. European Journal of Neuroscience 59, 1099–1140 (2024).

7 Lutas, A. et al. State-specific gating of salient cues by midbrain dopaminergic input to basal amygdala. Nature Neuroscience 22, 1820–1833 (2019).

8 Tang, W., Kochubey, O., Kintscher, M. & Schneggenburger, R. A VTA to Basal Amygdala Dopamine Projection Contributes to Signal Salient Somatosensory Events during Fear Learning. The Journal of Neuroscience 40, 3969 (2020).

9 Nader, K. & LeDoux, J. Inhibition of the mesoamygdala dopaminergic pathway impairs the retrieval of conditioned fear associations. Behavioral Neuroscience 113, 891–901 (1999).

10 Wilkinson, L. S. et al. Dissociations in dopamine release in medial prefrontal cortex and ventral striatum during the acquisition and extinction of classical aversive conditioning in the rat. European Journal of Neuroscience 10, 1019–1026 (1998).

11 Yan, R., Wang, T. & Zhou, Q. Elevated dopamine signaling from ventral tegmental area to prefrontal cortical parvalbumin neurons drives conditioned inhibition. Proceedings of the National Academy of Sciences 116, 13077–13086 (2019).

12 Menegas, W., Akiti, K., Amo, R., Uchida, N. & Watabe-Uchida, M. Dopamine neurons projecting to the posterior striatum reinforce avoidance of threatening stimuli. Nature Neuroscience 21, 1421–1430 (2018).

13 de Jong, J. W. et al. A Neural Circuit Mechanism for Encoding Aversive Stimuli in the Mesolimbic Dopamine System. Neuron 101, 133–151.e137 (2019).

14 Cai, L. X. et al. Distinct signals in medial and lateral VTA dopamine neurons modulate fear extinction at different times. eLife 9, e54936 (2020).

15 Chen, A. P. F. et al. Nigrostriatal dopamine modulates the striatal-amygdala pathway in auditory fear conditioning. Nature Communications 14, 7231 (2023).

16 Menegas, W. et al. Dopamine neurons projecting to the posterior striatum form an anatomically distinct subclass. eLife 4, e10032 (2015).

17 Menegas, W., Babayan, B. M., Uchida, N. & Watabe-Uchida, M. Opposite initialization to novel cues in dopamine signaling in ventral and posterior striatum in mice. eLife 6, e21886 (2017).

18 Green, I., Amo, R. & Watabe-Uchida, M. Shifting attention to orient or avoid: a unifying account of the tail of the striatum and its dopaminergic inputs. Current Opinion in Behavioral Sciences 59, 101441 (2024).

19 Kim, H. F., Ghazizadeh, A. & Hikosaka, O. Separate groups of dopamine neurons innervate caudate head and tail encoding flexible and stable value memories. Frontiers in Neuroanatomy 8 (2014).

20 Akiti, K. et al. Striatal dopamine explains novelty-induced behavioral dynamics and individual variability in threat prediction. Neuron 110, 3789–3804.e3789 (2022).

21 Tsutsui-Kimura, I., Uchida, N. & Watabe-Uchida, M. Dynamical management of potential threats regulated by dopamine and direct- and indirect-pathway neurons in the tail of the striatum. bioRxiv, 2022.2002.2005.479267 (2022).

22 Steinberg, E. E. et al. Amygdala-Midbrain Connections Modulate Appetitive and Aversive Learning. Neuron 106, 1026–1043.e1029 (2020).

23 LeDoux, J. E. Coming to terms with fear. Proceedings of the National Academy of Sciences 111, 2871–2878 (2014).

24 Romanski, L. M. & LeDoux, J. E. Information Cascade from Primary Auditory Cortex to the Amygdala: Corticocortical and Corticoamygdaloid Projections of Temporal Cortex in the Rat. Cerebral Cortex 3, 515–532 (1993).

25 Znamenskiy, P. & Zador, A. M. Corticostriatal neurons in auditory cortex drive decisions during auditory discrimination. Nature 497, 482–485 (2013).

26 Dalmay, T. et al. A Critical Role for Neocortical Processing of Threat Memory. Neuron 104, 1180–1194.e1187 (2019).

27 Watabe-Uchida, M., Zhu, L., Ogawa Sachie K., Vamanrao, A. & Uchida, N. Whole-Brain Mapping of Direct Inputs to Midbrain Dopamine Neurons. Neuron 74, 858–873 (2012).

28 Lerner, T. N. et al. Intact-Brain Analyses Reveal Distinct Information Carried by SNc Dopamine Subcircuits. Cell 162, 635–647 (2015).

29 Wu, J. et al. Distinct Connectivity and Functionality of Aldehyde Dehydrogenase 1a1-Positive Nigrostriatal Dopaminergic Neurons in Motor Learning. Cell Reports 28, 1167–1181.e1167 (2019).

30 Thompson, W. S., Grillner, S. & Silberberg, G. Motor cortex directly excites the output nucleus of the basal ganglia, the substantia nigra pars reticulata. bioRxiv, 2024.2003.2019.585632 (2024).

31 Fleming, L. L., Harnett, N. G. & Ressler, K. J. Sensory alterations in post-traumatic stress disorder. Current Opinion in Neurobiology 84, 102821 (2024).

32 Liang, C. L., Sinton, C. M. & German, D. C. Midbrain dopaminergic neurons in the mouse: co-localization with Calbindin-D28k and calretinin. Neuroscience 75, 523–533 (1996).

33 Yamaguchi, T., Wang, H. L. & Morales, M. Glutamate neurons in the substantia nigra compacta and retrorubral field. Eur J Neurosci 38, 3602–3610 (2013).

34 Poulin, J.-F. et al. Mapping projections of molecularly defined dopamine neuron subtypes using intersectional genetic approaches. Nature Neuroscience 21, 1260–1271 (2018).

35 Poulin, J. F., Gaertner, Z., Moreno-Ramos, O. A. & Awatramani, R. Classification of Midbrain Dopamine Neurons Using Single-Cell Gene Expression Profiling Approaches. Trends Neurosci 43, 155–169 (2020).

36 Nemoto, C., Hida, T. & Arai, R. Calretinin and calbindin-D28k in dopaminergic neurons of the rat midbrain: a triple-labeling immunohistochemical study. Brain Research 846, 129–136 (1999).

37 Kang, Y. & Kitai, S. T. A whole cell patch-clamp study on the pacemaker potential in dopaminergic neurons of rat substantia nigra compacta. Neuroscience Research 18, 209–221 (1993).

38 Tarfa, R. A., Evans, R. C. & Khaliq, Z. M. Sensitivity to Hyperpolarizing Inhibition in Mesoaccumbal Relative to Nigrostriatal Dopamine Neuron Subpopulations. The Journal of Neuroscience 37, 3311 (2017).

39 Henny, P. et al. Structural correlates of heterogeneous in vivo activity of midbrain dopaminergic neurons. Nature Neuroscience 15, 613–619 (2012).

40 Feigin, L., Tasaka, G., Maor, I. & Mizrahi, A. Sparse Coding in Temporal Association Cortex Improves Complex Sound Discriminability. The Journal of Neuroscience 41, 7048 (2021).

41 Galtieri, D. J., Estep, C. M., Wokosin, D. L., Traynelis, S. & Surmeier, D. J. Pedunculopontine glutamatergic neurons control spike patterning in substantia nigra dopaminergic neurons. eLife 6, e30352 (2017).

42 Smith, I. D. & Grace, A. A. Role of the subthalamic nucleus in the regulation of nigral dopamine neuron activity. Synapse 12, 287–303 (1992).

43 Iribe, Y., Moore, K., Pang, K. C. H. & Tepper, J. M. Subthalamic Stimulation-Induced Synaptic Responses in Substantia Nigra Pars Compacta Dopaminergic Neurons In Vitro. Journal of Neurophysiology 82, 925–933 (1999).

44 Futami, T., Takakusaki, K. & Kitai, S. T. Glutamatergic and cholinergic inputs from the pedunculopontine tegmental nucleus to dopamine neurons in the substantia nigra pars compacta. Neuroscience Research 21, 331–342 (1995).

45 Bingmin, L. et al. Frequency-Dependent Plasticity in the Temporal Association Cortex Originates from the Primary Auditory Cortex, and Is Modified by the Secondary Auditory Cortex and the Medial Geniculate Body. The Journal of Neuroscience 42, 5254 (2022).

46 Neuhoff, H., Neu, A., Liss, B. & Roeper, J. I(h) channels contribute to the different functional properties of identified dopaminergic subpopulations in the midbrain. J Neurosci 22, 1290–1302 (2002).

47 Farassat, N. et al. In vivo functional diversity of midbrain dopamine neurons within identified axonal projections. eLife 8, e48408 (2019).

48 Evans, R. C., Zhu, M. & Khaliq, Z. M. Dopamine Inhibition Differentially Controls Excitability of Substantia Nigra Dopamine Neuron Subpopulations through T-Type Calcium Channels. The Journal of Neuroscience 37, 3704 (2017).

49 Evans, R. C. et al. Functional Dissection of Basal Ganglia Inhibitory Inputs onto Substantia Nigra Dopaminergic Neurons. Cell Reports 32, 108156 (2020).

50 Tucker, K. R., Huertas, M. A., Horn, J. P., Canavier, C. C. & Levitan, E. S. Pacemaker Rate and Depolarization Block in Nigral Dopamine Neurons: A Somatic Sodium Channel Balancing Act. The Journal of Neuroscience 32, 14519 (2012).

51 Dodson, P. D. et al. Representation of spontaneous movement by dopaminergic neurons is cell-type selective and disrupted in parkinsonism. Proceedings of the National Academy of Sciences 113, E2180–E2188 (2016).

52 Cobb-Lewis, D. E., Sansalone, L. & Khaliq, Z. M. Contributions of the Sodium Leak Channel NALCN to Pacemaking of Medial Ventral Tegmental Area and Substantia Nigra Dopaminergic Neurons. The Journal of Neuroscience 43, 6841 (2023).

53 Yang, H. et al. Nucleus Accumbens Subnuclei Regulate Motivated Behavior via Direct Inhibition and Disinhibition of VTA Dopamine Subpopulations. Neuron 97, 434–449.e434 (2018).

54 Kramer, D. J., Risso, D., Kosillo, P., Ngai, J. & Bateup, H. S. Combinatorial Expression of Grp and Neurod6 Defines Dopamine Neuron Populations with Distinct Projection Patterns and Disease Vulnerability. eneuro 5, ENEURO.0152-0118.2018 (2018).

55 Todd, K. L., Lipski, J. & Freestone, P. S. Subthalamic nucleus exclusively evokes dopamine release in the tail of the striatum. Journal of Neurochemistry 162, 417–429 (2022).

56 Bruet, N. et al. High Frequency Stimulation of the Subthalamic Nucleus Increases the Extracellular Contents of Striatal Dopamine in Normal and Partially Dopaminergic Denervated Rats. Journal of Neuropathology & Experimental Neurology 60, 15–24 (2001).

57 Shon, Y.-M. et al. High frequency stimulation of the subthalamic nucleus evokes striatal dopamine release in a large animal model of human DBS neurosurgery. Neuroscience Letters 475, 136–140 (2010).

58 Frankle, W. G., Laruelle, M. & Haber, S. N. Prefrontal Cortical Projections to the Midbrain in Primates: Evidence for a Sparse Connection. Neuropsychopharmacology 31, 1627–1636 (2006).

59 Cho, J.-H., Huang, B. S. & Gray, J. M. RNA sequencing from neural ensembles activated during fear conditioning in the mouse temporal association cortex. Scientific Reports 6, 31753 (2016).

60 Matsushita, N. et al. Dynamics of tyrosine hydroxylase promoter activity during midbrain dopaminergic neuron development. Journal of Neurochemistry 82, 295–304 (2002).

