## Supplementary material for "Corticonigral projections recruit substantia nigra pars lateralis dopaminergic neurons for auditory threat memories": Sansalone et al 2024_Supplementary Material

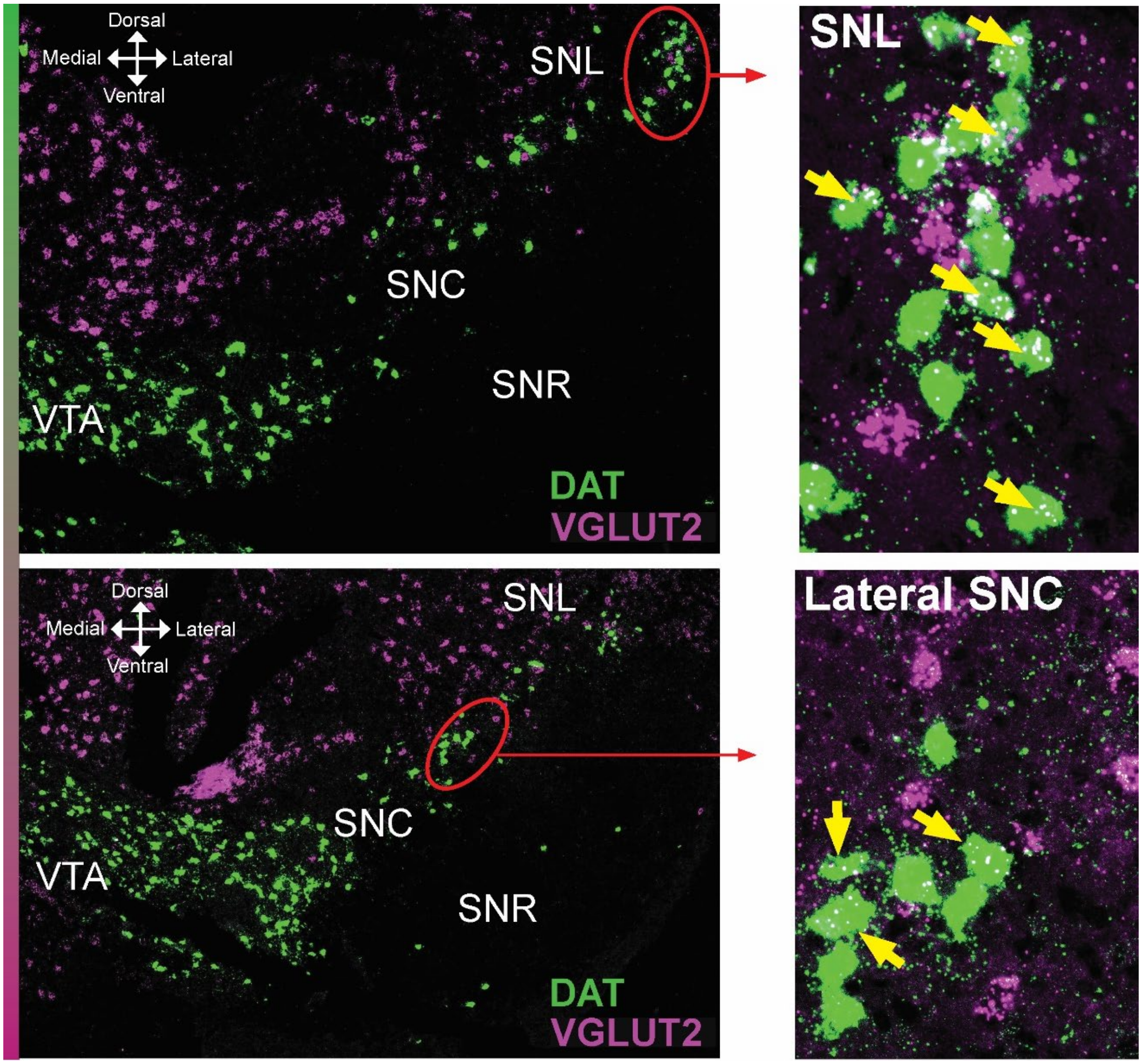


**Extended Figure 1 I Left,** In situ hybridization experiment showing a coronal section of substantia nigra from a C57WT mouse with DAT+ (green) and VGluT2+ (purple) neurons. **Right,** Yellow arrows indicate doubly positive neurons for DAT and VGluT2 in SNL (top) and lateral SNC (bottom).

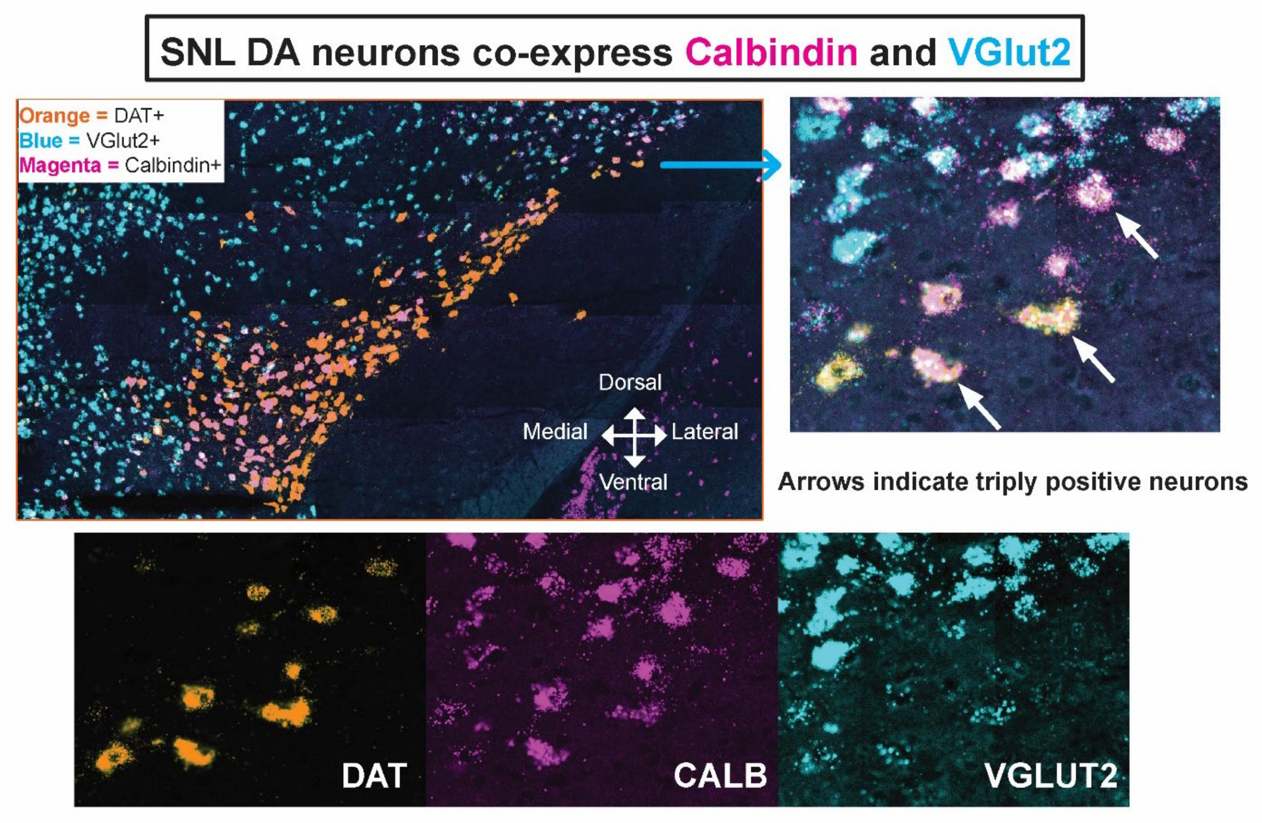


**Extended Figure 2 I Top left,** In situ hybridization experiment showing a coronal section from a C57WT mouse with DAT+ (orange), Calbindin+ (magenta) and VGluT2+ (blue) neurons. **Top right**, Magnification of the SNL showing triply positive neurons for DAT, Calb and VGluT2. **Bottom,** Split channels for top right image.

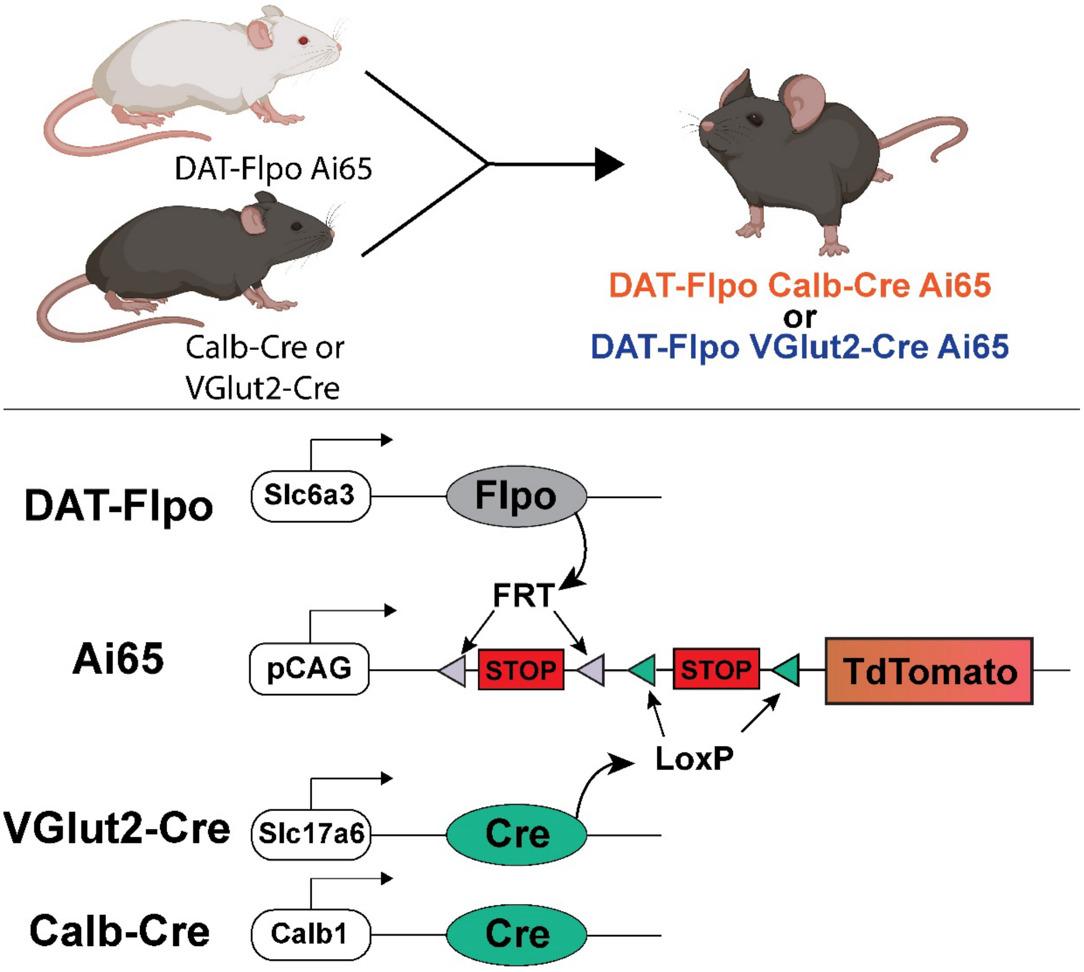


**Extended Figure 3 I a,b** Intersectional genetic strategy used to generate DAT-Flp Calb-Cre or DAT-Flp VGluT2-Cre mice.

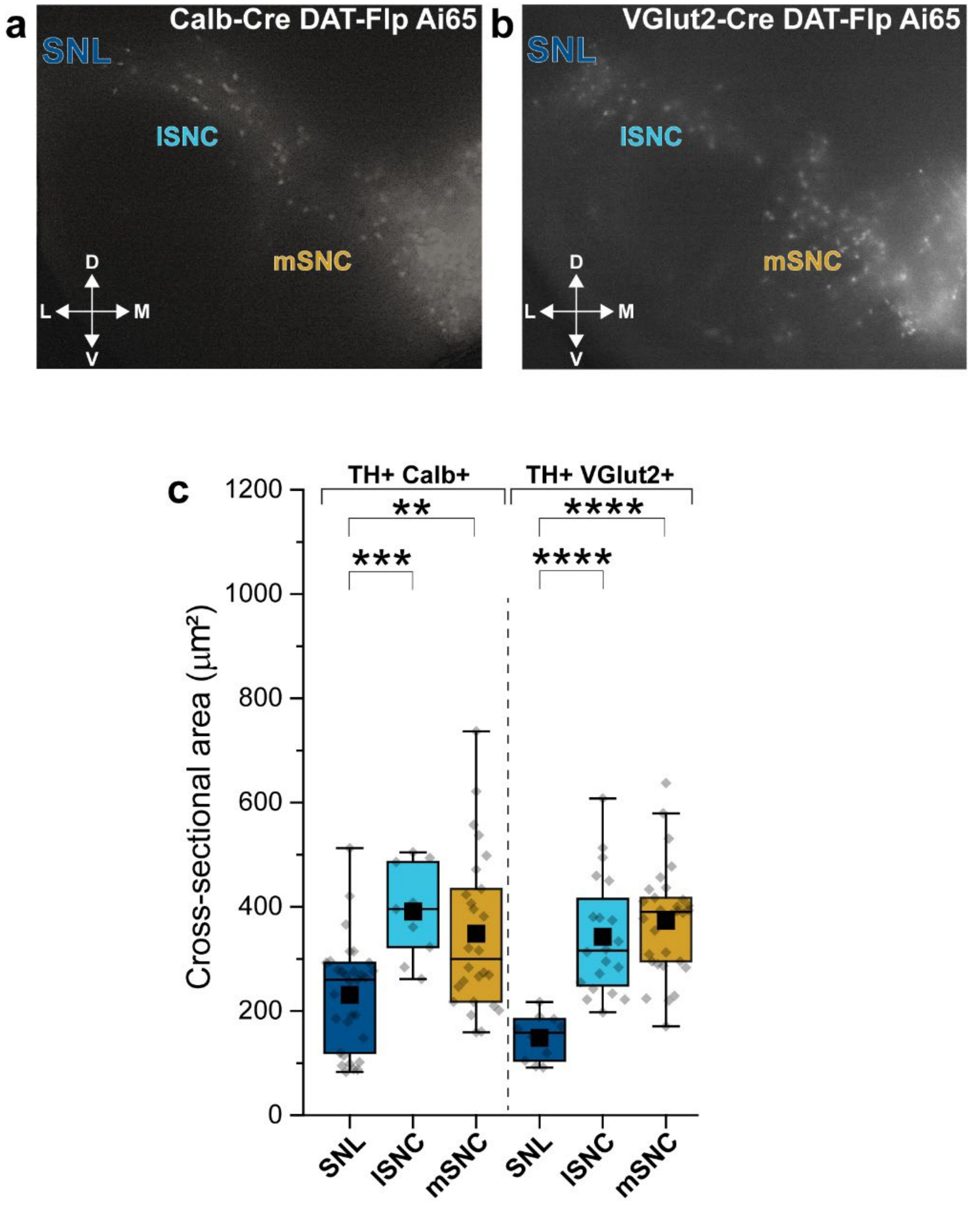


**Extended Figure 4 I a,b** Representative coronal sections from Calb-Cre DAT-Flp Ai65 or VGluT2-Cre DAT-Flp Ai65 mice showing Calb+ (**a**) or VGluT2+ (**b**) DANs (white) along the mediolateral axis of the substantia nigra. **c.** Bar plots showing cross-sectional somatic areas from TH+/Calb+ or TH+/VGluT2+ neurons with means (All compared with SNL, Mann-Whitney test).

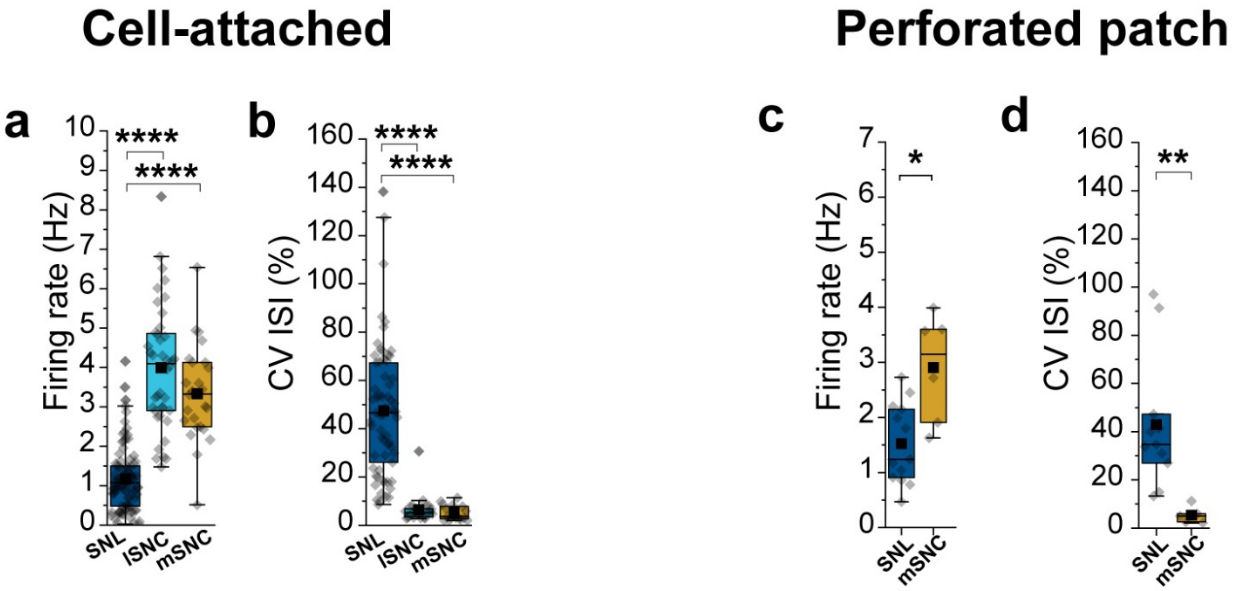


**Extended Figure 5 I** Bar plots with means showing firing rate and CV ISI of DANs obtained from electrophysiology recordings in brain slices from DAT-Cre Ai9, Calb-Cre DAT-Flp Ai65 and VGluT2-Cre DAT-Flp Ai65 mice. **a,** Cell-attached Firing rate; SNL, n = 90, 1.17 ± 0.09 Hz; mSNC, n = 26, 3.33 ± 0.24 Hz; lSNc, n = 40, 3.99 ± 0.25 Hz; SNL vs mSNC, p = 1.53 x $10^{-14}$, SNL vs lSNC, p = 8.74 x $10^{-17}$). **b,** Cell-attached CV ISI; SNL, n = 65, 47.44 ± 3.39 %; mSNC, n = 17, 5.28 ± 0.75 %; lSNc, n = 25, 6.42 ± 1.08 %; SNL vs mSNC, p = 2.35 x 10-$^{16}$, SNL vs lSNC, p = 7.76 x $10^{-19}$). **c,** Perforated-patch Firing rate; SNL, n = 13, 1.52 ± 0.20 Hz; mSNC, n = 6, 2.90 ± 0.40 Hz; SNL vs mSNC, p = 0.017). **d,** Perforated-patch CV ISI; SNL, n = 13, 55.18 ± 13.55 %; mSNC, n = 6, 11.44 ± 6.23 %; SNL vs mSNC, p = 0.005).

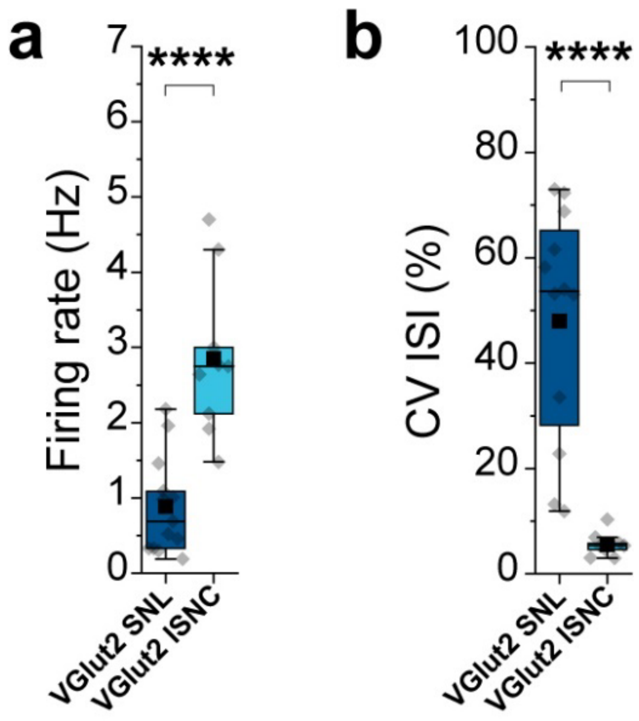


**Extended Figure 6 I** Bar plots with means showing firing rate and CV ISI of DANs obtained from whole-cell recordings in brain slices from VGluT2-Cre DAT-Flp Ai65 mice. **a,** Cell-attached Firing rate (SNL, n = 13, 0.88 ± 0.18 Hz; lSNc, n = 9, 2.85 ± 0.35 Hz; SNL vs lSNC, $p = 7.64 \times 10^{-5}$). **b,** Cell-attached CV ISI (SNL, n = 12, 47.96 ± 6.39%; lSNc, n = 9, 5.55 ± 0.73 %; SNL vs lSNC, $p = 6.80 \times 10^{-6}$).

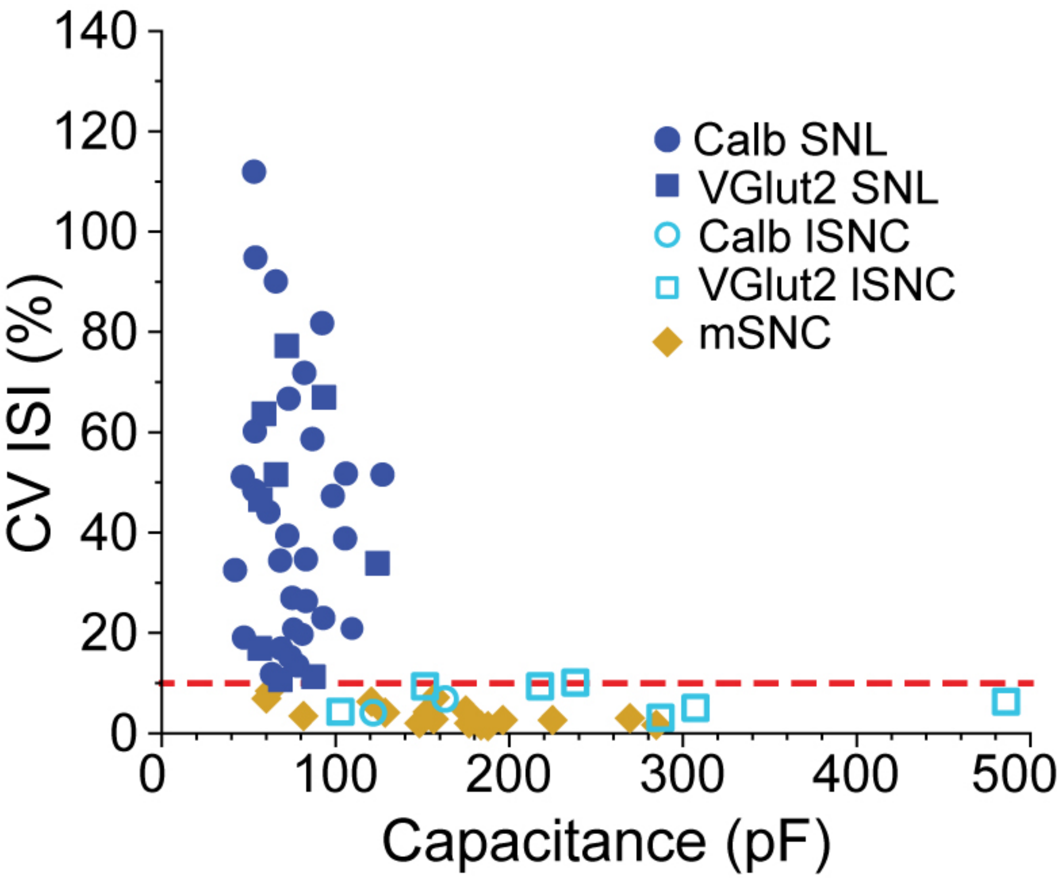


**Extended Figure 7 I** Scatter graph showing correlation between coefficient of variation of interspike interval (CV ISI) and capacitance (pF). Data obtained from whole-cell patch clamp recordings in DANs from Calb-Cre DAT-Flp Ai65, VGluT2-Cre DAT-Flp Ai65, DAT-Cre Ai9 mice and TH-GFP mice. Dotted red showing that, virtually, all SNc DANs have a CV ISI equal or lower than 10%.

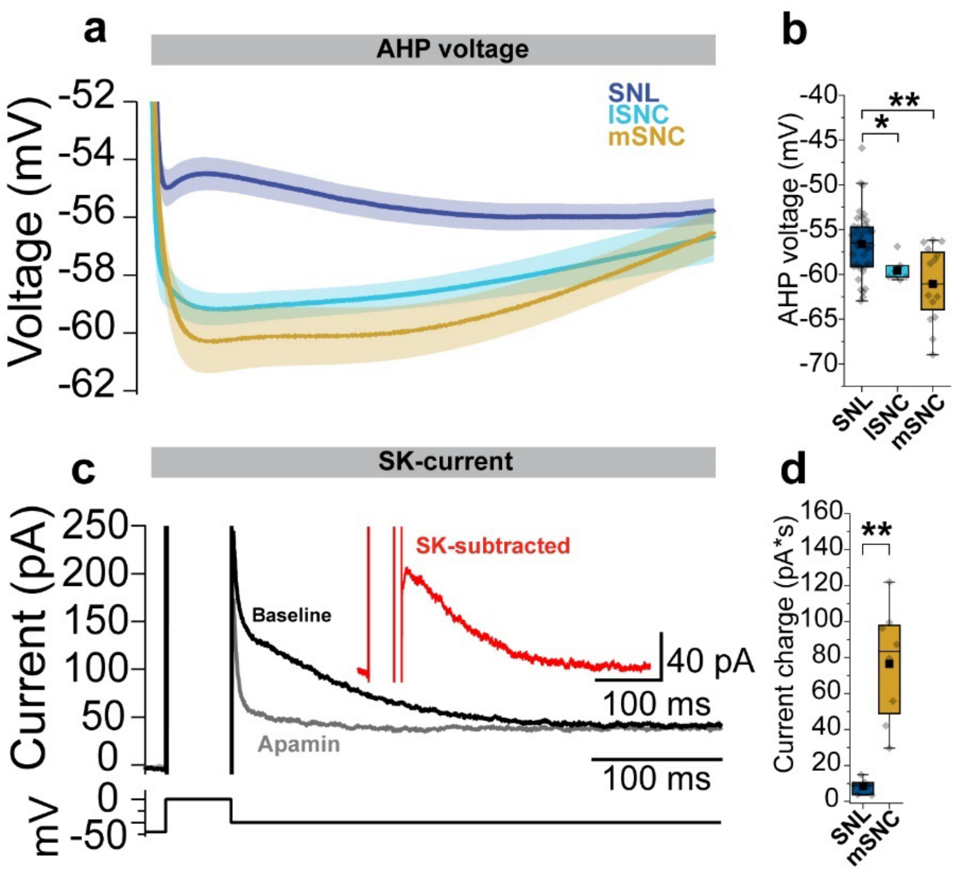


**Extended Figure 8 I a,** Comparison of AHP voltage during whole-cell patch-clamp recordings in SNL (blue), lSNc (cyan) and mSNc (ochre) DANs from DAT-Cre Ai9 and TH-GFP mice. **b**, Bar plots showing that average AHP voltage minima for SNL DANs are significantly smaller compared to lSNc and mSNc DANs (Average AHP minimum; SNL, n = 41, -56.61 ± 0.55 mV; lSNc, n = 7, -59.58 ± 0.49 mV; mSNc, n = 16, 61.07 ±1.00 mV; SNL vs lSNc, p = 0.012; SNL vs lSNc, p = 4.62 x $10^{-4}$). **c,** The AHP is significantly affected by small-conductance calcium-activated potassium currents (SK). A more depolarized AHP for SNL DANs suggests they have lower levels of SK channels. Thus, we performed whole-cell voltage clamp recordings to estimate SK currents in SNL and SNc DANs from DAT-Cre Ai9 and TH-GFP mice. **d,** Bar plots showing that average current charges for SNL are significantly lower compared to mSNc DANs (Current charge; SNL, n = 6, 8.42 ± 1.76 pA*S; mSNc, n = 8, 76.5 ±11.13 pA*S; p = 0.0022).

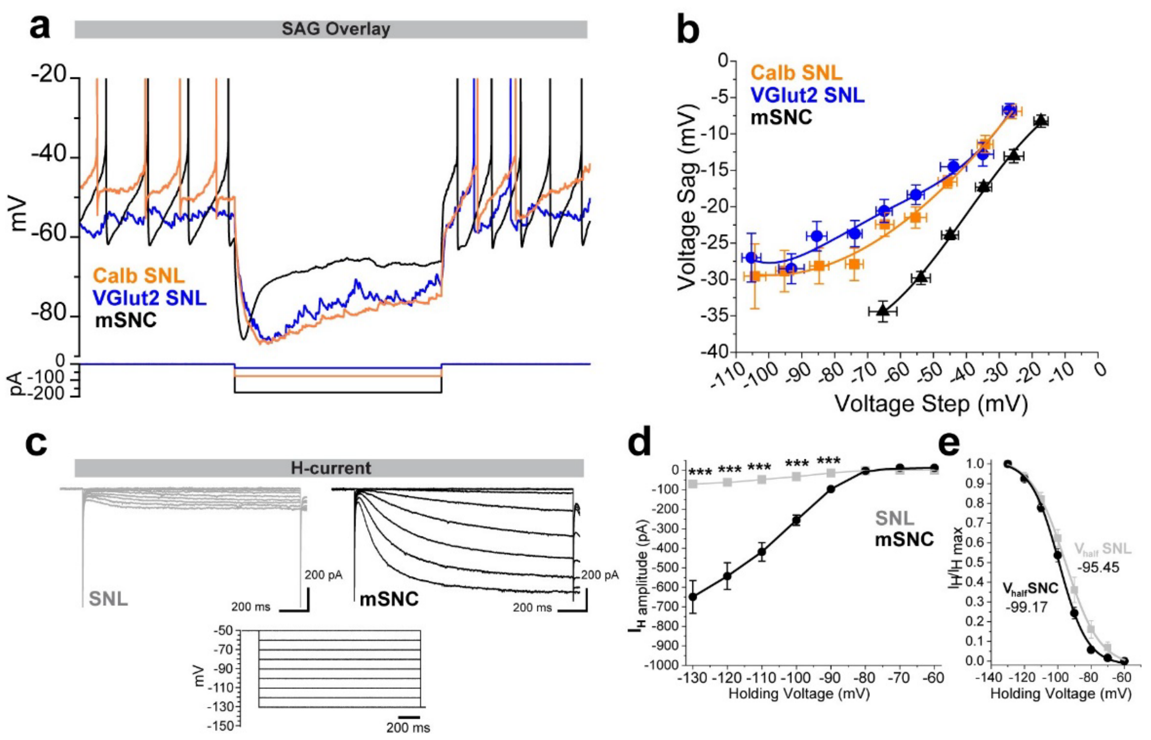


**Extended Figure 9 I a,** Whole-cell recordings showing SAG overlay for SNL (orange, blue) and mSNC (black) DANs from Calb-Cre DAT-Flp Ai65, VGluT2-Cre DAT-Flp Ai65 and DAT-Cre Ai9 mice. **b,** Voltage dependence of SAG in SNL and mSNc DANs from same mice in **a**. **c,** DANs SAG is mainly generated by hyperpolarization-activated cyclic nucleotide-gated channels (HCN). Here, we isolated HCN conductances at decreasing voltages (bottom) in SNL (top left, grey) and mSNc (top right, black) DANs from VGluT2-Cre DAT-Flp Ai65 and DAT-Cre Ai9 mice. **d,** Voltage dependence of HCN current amplitude ($I_H$) showing that SNL DANs display significantly lower HCN currents compared to mSNc DANs (max current amplitude; SNL, n = 8, - 70.8 ± 11.60 pA; mSNc, n = 7, - 649.07 ± 84.09; $p = 3.1 \times 10^{-4}$). **e,** Plot showing $I_H$ activation voltage showing similar profiles for SNL and mSNc DANs.

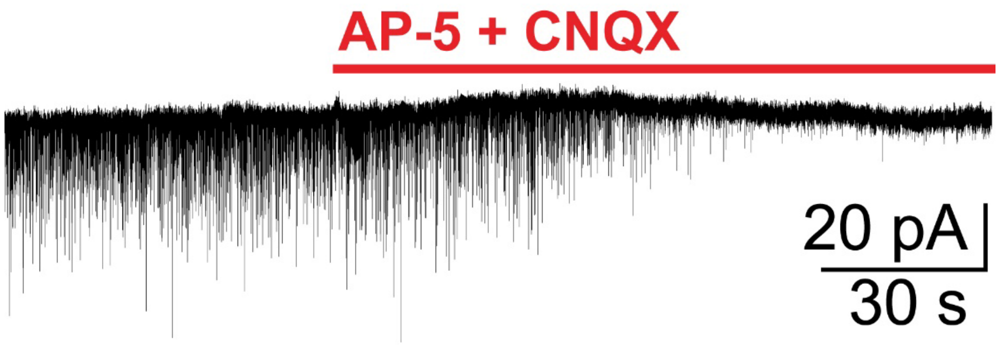


**Extended Figure 10 I** Voltage clamp recording in SNL DAN from a VGluT2-Cre mouse. Neuron was held at – 70 mV. Note the suppression of virtually all post-synaptic currents (PSCs) upon bath application of D-AP5 (50 μM) and CNQX (μM).

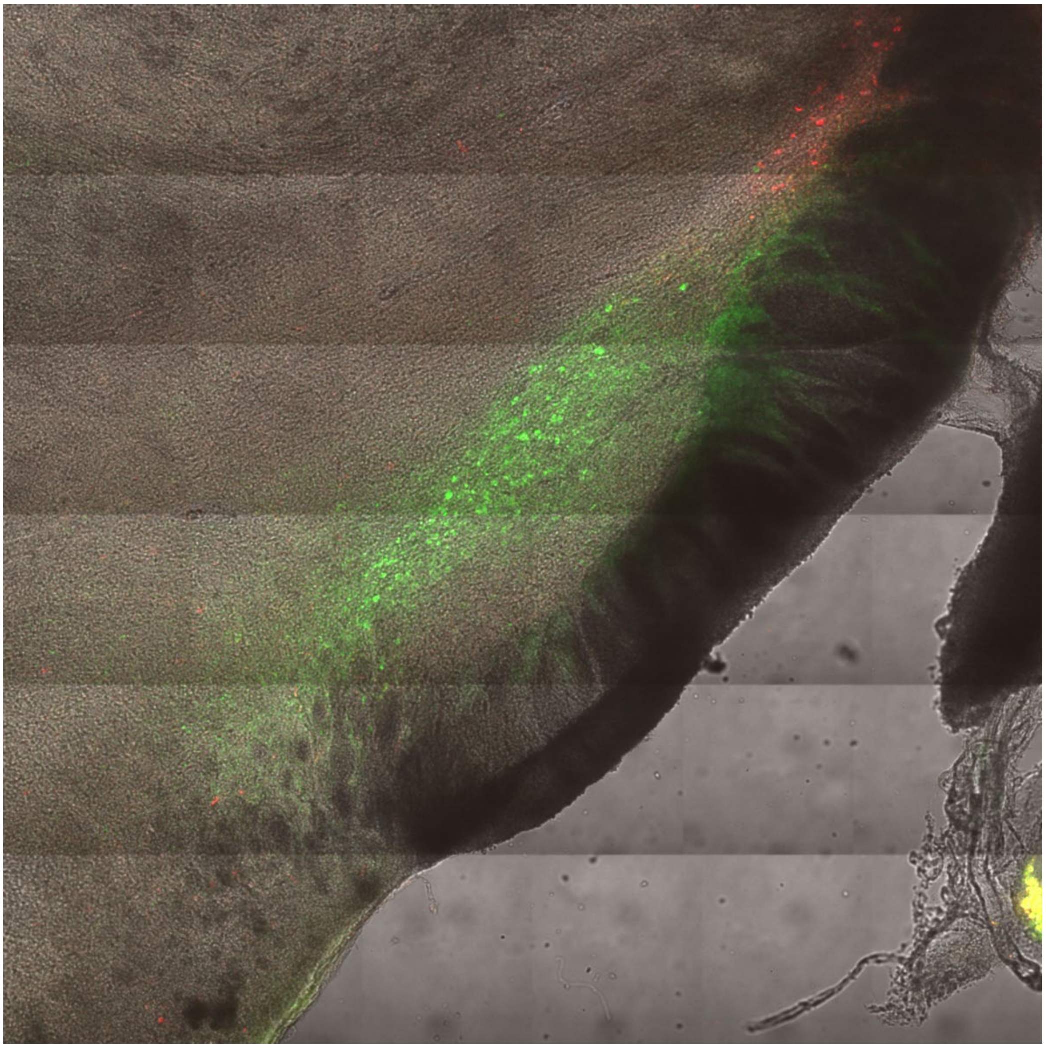

**Extended data figure 11 I** Confocal image showing coronal slice with STN from C57WT mouse injected with CTB55 in lSNc and CTB647 in SNL. Note that most retrogradely labeled neurons from lSNc are in the central section of the STN while retrogradely labeled neurons from SNL are located dorsolaterally.

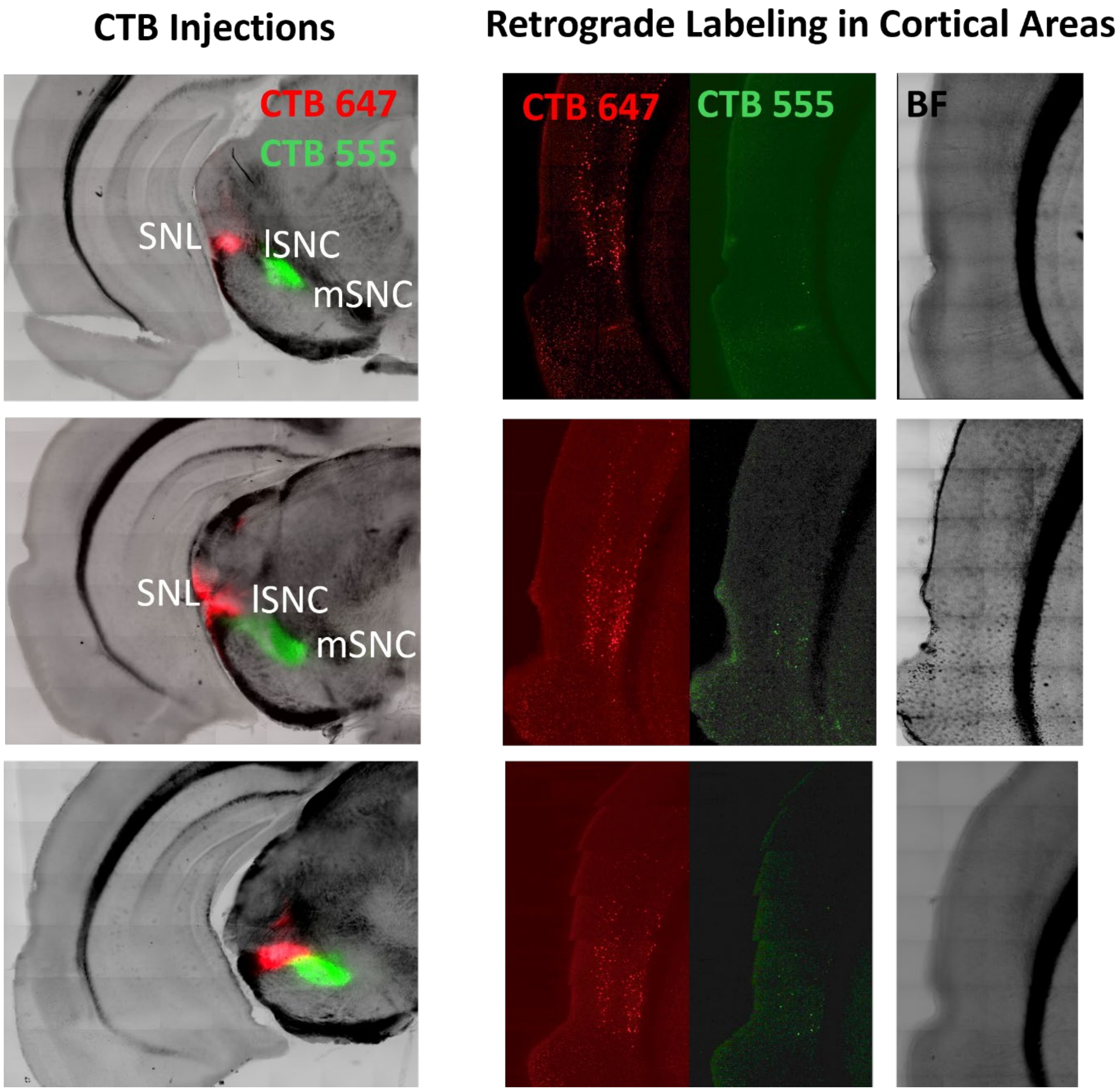


**Extended data figure 12 I Left**, confocal images showing progressive coronal sections from a C57WT mouse that was injected in SNL and lSNc with CTB555 (green) and CTB647 (red), respectively. **Right**, confocal images showing retrogradely labeled neurons in layers 2/3 and 5 of auditory cortex (EcT/TeA/AuV) from SNL injections (red). Note that retrograde labeling from lSNc injection (green) is virtually absent, and the sparse neurons shown in green likely represent labeling through viral spread of CTB555 injection into SNL.
